# Benchmarking 16S rRNA gene amplicon analysis in high-diversity microbial communities reveals fundamental trade-offs in clustering and denoising

**DOI:** 10.64898/2026.09.21.752605

**Authors:** Cassandra Stamsaas, Torbjørn Rognes, Knut Rudi, Lars Snipen, Hilde Vinje

## Abstract

**Background:** Amplicon sequencing of the 16S rRNA gene is widely used to characterise microbial communities, but the performance of commonly used clustering and denoising pipelines has not been systematically evaluated for highly diverse environmental datasets. We therefore benchmarked four established clustering and denoising pipelines, VSEARCH *cluster_size*, UNOISE as implemented in VSEARCH, Swarm, and DADA2, using simulated microbial communities with known ground-truth compositions and evaluated how performance varied with species richness, abundance unevenness, sequencing depth, and minimum abundance threshold.

**Results:** Differences among pipelines were small at low species richness but became greater as species richness increased. High average cluster purity did not necessarily correspond to accurate species-level recovery, as species could be split across multiple clusters or recovered incompletely. Among the pipelines, UNOISE may be preferable when high species representation and cluster purity are prioritised, while also showing strong reconstruction of relative abundance profiles, albeit with extensive species splitting and many unclustered reads. DADA2 showed similarly strong reconstruction of relative abundance profiles and less species splitting but represented fewer species and had lower cluster purity at higher richness. Swarm provided a balanced compromise between species representation and splitting, whereas *cluster_size* may be useful when limiting species splitting is a priority, despite weaker reconstruction of relative abundance profiles. Increasing the minimum abundance threshold reduced species splitting but also reduced the number of clusters and perfect clusters and, at higher thresholds, decreased concordance with ground-truth relative abundance profiles. These effects were more pronounced at lower sequencing depths. Analysis of seafloor sediment samples also showed threshold-dependent sequence loss, including the loss of sequences repeatedly detected across replicate samples.

**Conclusions:** Pipeline performance in highly diverse 16S rRNA amplicon datasets varies with dataset characteristics, evaluation criteria, and parameter settings. Pipeline selection should therefore reflect dataset characteristics and the analytical objective rather than rely on a single measure of performance, and minimum abundance thresholds should be evaluated in relation to sequencing depth rather than applied as fixed defaults. These findings emphasise the importance of reconsidering established analytical practices and transparently reporting bioinformatic settings as 16S rRNA amplicon sequencing is increasingly applied to highly diverse environmental microbial communities.

## Background

Metabarcoding has become a central approach for characterising biodiversity, community composition, and ecological change across diverse environments [1]. Among metabarcoding approaches, amplicon sequencing of 16S rRNA genes is widely used to characterise microbial communities [2], [3], offering a cost-effective and scalable method that enables deep sequencing across large numbers of samples. While 16S rRNA metabarcoding has been extensively studied and validated in relatively well-characterised host-associated microbiomes [4], such as the gut, it is increasingly being applied to highly diverse environmental microbial communities in soils, sediments, and aquatic environments.

The rapidly expanding use of environmental DNA (eDNA)-based approaches to address ecological and environmental questions has extended microbial community profiling to increasingly diverse and complex environments. Samples from outdoor environments, such as soil, sediments and aquatic systems, often contain microbial communities characterised by high species richness, uneven abundance distributions, and potentially many closely related taxa [5], [6], [7], [8]. In such communities, the likelihood of encountering closely related taxa with similar amplicon sequences increases, making it more challenging to distinguish true biological variation from technical noise.

However, the development and evaluation of bioinformatic methods have not kept pace with this shift towards increasingly diverse environmental applications. Many commonly used computational pipelines were originally developed and evaluated primarily using mock communities or comparatively well-characterised microbiome datasets, whereas highly diverse environmental samples remain considerably more difficult to benchmark [9], [10], [11]. The increased richness, sequence similarity, and uneven abundance distributions characteristic of environmental datasets present substantially greater challenges for denoising and clustering than those encountered in low-diversity systems. Consequently, analytical choices that have only minor effects in low-diversity systems may have substantially larger impacts in complex environmental communities. These effects may influence downstream estimates of diversity and community composition and, ultimately, the biological conclusions drawn from these studies.

Despite the widespread use of bioinformatics pipelines, relatively few benchmarking studies have systematically assessed how these pipelines perform under conditions resembling highly diverse environmental communities [10], [12], [13], [14], [15]. A central obstacle is the lack of reliable ground truth in real environmental datasets, which limits direct evaluation of accuracy, error rates, and recovery of the underlying community. Previous benchmarking studies have partly addressed this challenge using indirect evaluation strategies, including consistency across technical replicates and prior biological knowledge about expected diversity patterns [14]. However, such approaches do not provide fully specified read- or species-level ground truth. Although well-defined mock communities provide useful frameworks for validation, their limited richness and taxonomic complexity can constrain their relevance for benchmarking methods intended for heterogeneous environmental datasets [16].

Simulation studies offer a direct solution to this limitation. Because simulated reads are generated from predefined community compositions, they enable systematic evaluation against known ground truth, including assessment of how pipelines respond to increasing diversity and where deviations from the expected community structure arise [17], [18]. While pipeline choice is often emphasised in metabarcoding workflows, outcomes are strongly influenced by user-defined parameter settings. This issue may be particularly important in highly diverse environmental communities, where distinguishing biological variants from sequencing artefacts is inherently challenging. However, default parameter values are often applied without systematic evaluation in relation to dataset-specific properties, despite evidence that analytical settings can substantially influence results [9], [10]. Simulated datasets also provide an opportunity to address this limitation by enabling direct assessment of how user-defined parameter settings influence clustering and denoising performance across communities of varying complexity. Among these settings, the minimum abundance threshold is particularly relevant in several pipelines, as it determines which dereplicated sequences are retained for downstream clustering or denoising. This parameter affects the balance between removing likely errors and retaining rare, potentially genuine biological variants.

Guidance on how to set the minimum abundance threshold remains limited, and systematic evaluations across datasets of varying complexity are lacking. Available recommendations are often restricted to default settings or brief comments in pipeline documentation [19]. This represents an important methodological gap, particularly for high-diversity datasets, in which the distinction between rare biological variants and technical artefacts is especially challenging.

To address these methodological gaps, we systematically investigated how clustering and denoising pipelines respond to increasing species richness and maximum-to-minimum abundance ratio. We evaluated four established pipelines: VSEARCH (*cluster_size*) [20], UNOISE [21] (as implemented in VSEARCH [20]), Swarm [22], and DADA2 [23]. We used simulated datasets based on predefined community structures with known ground-truth across gradients of species richness, maximum-to-minimum abundance ratio, and sequencing depth. Pipeline performance was assessed using ground-truth-based metrics of cluster quality, recovery of simulated taxa, and reconstruction of true abundance profiles. We also examined how the minimum abundance threshold influences performance across datasets of varying diversity, with the aim of assessing whether default settings are suitable for high-diversity microbial community analyses. Finally, to examine whether patterns observed in simulations were reflected in environmental samples, we applied the same pipelines to seafloor sediment data as a complementary empirical case study. Although these data lack fully specified ground truth, they provide an opportunity to assess whether trends observed under controlled simulation conditions are also evident in highly diverse natural microbial communities.

## Materials and methods

We generated six simulated community configurations representing Illumina 16S rRNA amplicon sequencing data. The configurations were defined by three levels of species richness and two maximum-to-minimum abundance ratios. Each configuration was simulated at three sequencing depths, with 20 replicate datasets generated at each sequencing depth. The resulting datasets were used to evaluate four established clustering and denoising pipelines: VSEARCH *cluster_size*, UNOISE (as implemented in VSEARCH), Swarm, and DADA2. The overall study design and benchmarking workflow are summarised in Figure 1.

**Figure 1.**
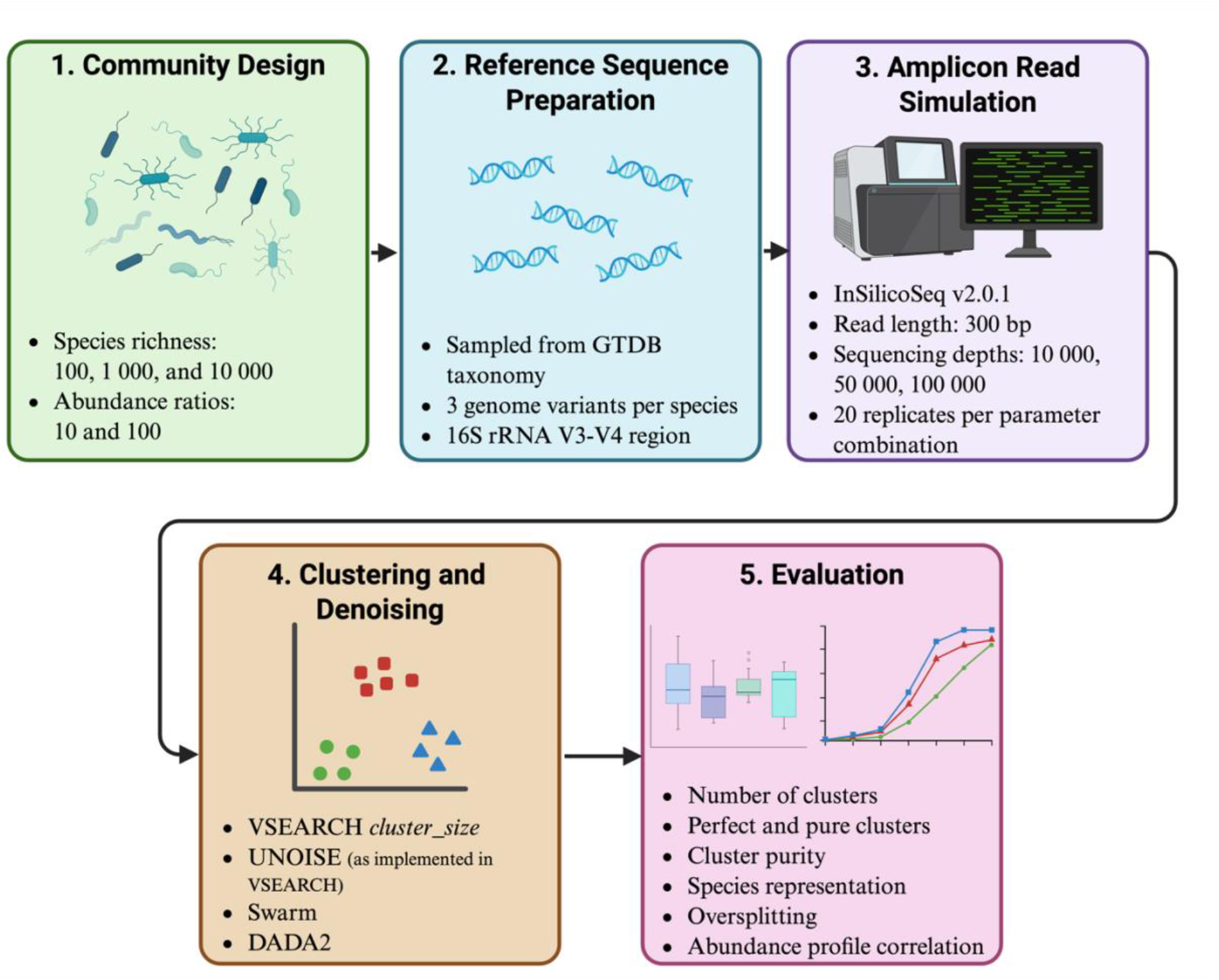
Overview of the study design and benchmarking workflow for generating and evaluating simulated 16S rRNA amplicon datasets. Microbial communities varying in species richness (100, 1 000, and 10 000 species) and maximum-to-minimum abundance ratios (10 and 100) were generated from GTDB-based reference sequences. Amplicon reads were simulated at three sequencing depths (10 000, 50 000, and 100 000) and analysed using VSEARCH cluster_size, UNOISE (as implemented in VSEARCH), Swarm, and DADA2. Performance was evaluated against the known ground truth using metrics of cluster quality, species representation, reconstruction of relative abundance profiles, and sequence loss. Created in BioRender. Stamsaas, C. (2027) https://BioRender.com/tn6xp49.

All R analyses were conducted using R version 4.5.1 [24]. Command-line tools were executed on an Apple Silicon (ARM64) platform or, for selected analyses, on the Orion High Performance Computing Center (OHPCC) at the Norwegian University of Life Sciences (NMBU). Downstream data processing and visualisation were performed in R. The code and scripts used to generate simulated datasets, run the analysis pipelines, and calculate benchmarking metrics, are available on GitHub (https://github.com/CassandraHjo/benchmarking-supplementary-scripts).

### Sequence simulation

To enable direct comparison with known ground truth, we generated simulated amplicon datasets in which the taxonomic origin and abundance of each read were known. In this study, community complexity was operationally characterised by two dimensions: species richness and abundance unevenness, represented by the maximum-to-minimum abundance ratio. Higher species richness increased the number of taxa to be reconstructed, whereas larger abundance ratios produced more uneven communities in which low-abundance taxa were more difficult to recover correctly. To vary these properties systematically and represent variation in environmental community complexity, we generated datasets spanning three levels of species richness (100, 1 000, and 10 000 species) and two maximum-to-minimum abundance ratios (10 and 100). Species abundances were sampled from an exponential distribution, and the maximum-to-minimum abundance ratio was defined as the abundance of the most abundant species divided by that of the least abundant species. For brevity, this metric is hereafter referred to as the abundance ratio.

Six simulated community configurations were generated using InSilicoSeq 2.0.1 [25], which simulates sequencing reads from a specified error model, a set of reference sequences, and associated metadata, including read counts. All simulations used the prebuilt MiSeq error model and a read length of 300 base pairs (bp). Reads were generated for each configuration described above at sequencing depths of 10 000, 50 000, and 100 000 reads, with 20 replicate datasets generated per configuration. Unless otherwise stated, analyses were based on datasets with a sequencing depth of 100 000 reads.

Species were randomly sampled without replacement from the unique species represented in a reference sequence dataset compiled by Vinje et al. [26], derived from the Genome Taxonomy Data Base project (GTDB) [27] release 2.26 (April 16, 2025). This dataset comprised 791 175 high-quality 16S rRNA sequences from the GTDB genomes, with multiple genomes and sequence variants represented for many species. For each selected species, up to three unique genomes were randomly sampled, and all available 16S rRNA variants associated with these genomes were retained. This allowed the simulated communities to include both variation among genomes within species and variation among 16S rRNA gene copies within individual genomes. Species abundances were assigned according to the exponential distribution specified by the selected abundance ratio and were then distributed uniformly among all sequences associated with each species. For each configuration, replicate datasets were generated by resampling species and independently simulating reads.

For each reference sequence, the 16S rRNA V3-V4 region was extracted using the primer pair Uni340F (5′-CCTACGGGRBGCASCAG-3′) and Uni806R (5′-GGACTACNNGGGTATCTAAT-3′), as described by Takai and Horikoshi [28]. Primer matching was performed in R using exact matching, and sequences lacking a match to the primer pair were discarded. The resulting amplicon sequences were used as reference sequences for read simulation with InSilicoSeq.

### Preprocessing of sequencing reads

To minimise differences attributable to read trimming and quality filtering, all datasets were subjected to the same initial trimming and filtering procedure before pipeline-specific downstream processing. Read trimming and quality filtering were performed using the *vs_fastx_trim_filt* function in Rsearch [29], with the minimum read length (*minlen*) set to 20 and the maximum expected error rate (*maxee_rate*) set to 0.01. The average expected error-per-base threshold was determined separately for each dataset by optimising the *truncee_rate* value using the *vs_optimize_truncee_rate* function in Rsearch [29] with default settings. The optimal truncation setting was defined as the value that maximised the number of merged read pairs with a copy number greater than two after dereplication.

The trimmed and filtered FASTQ files were used as input to the DADA2 pipeline, in which read merging and chimera removal were performed as part of the denoising process. For the VSEARCH-based pipelines and Swarm, reads were first merged and dereplicated, after which chimeric sequences were removed using the *vs_fastq_mergepairs*, *vs_fastx_uniques*, and *vs_uchime_denovo* functions in Rsearch. The resulting FASTA files were used as input for VSEARCH-based clustering and denoising and for Swarm.

### Clustering and denoising algorithms

We compared four established clustering and denoising pipelines: VSEARCH *cluster_size*, UNOISE [21] (as implemented in VSEARCH [20]), Swarm [22] and DADA2 [23]. In VSEARCH, we used the sequence similarity-based clustering algorithm *cluster_size* and the denoising algorithm UNOISE. For DADA2, we used the standard denoising pipeline, including error-rate learning (*learnErrors*), dereplication (*derepFastq*), denoising (*dada*), read merging (*mergePairs*), chimera removal (*removeBimeraDenovo*), and sequence-table construction (*makeSequenceTable*).

All VSEARCH commands were executed in R using the Rsearch package (v 1.2.0) [29] with VSEARCH 2.30.0 [20]. For *cluster_size*, dereplicated sequences with a copy number greater than or equal to the specified minimum abundance threshold were clustered at 99% sequence identity, whereas sequences below this threshold were subsequently mapped to the resulting centroids. The 99% sequence identity threshold was chosen because previous work by Vinje et al. [26] indicated that, within the GTDB taxonomy [27], clustering at 99% identity provides a near-optimal approximation of species-level groupings. The initial minimum abundance threshold was set to 2. Mapping was performed using global alignment (*vs_usearch_global*) with a 99% identity threshold. Sequences matching a centroid were assigned to the corresponding cluster, whereas sequences without a match were classified as unclustered and excluded from downstream cluster-based abundance estimates.

For denoising with UNOISE, two values of the *minsize* parameter were evaluated in the main pipeline comparison: 2 and 8. A *minsize* of 8 represents the default setting, whereas 2 was included to enable comparison with pipelines using a minimum abundance threshold of 2. For clarity, these two configurations are hereafter referred to as UNOISE 2 and UNOISE 8, respectively.

DADA2 version 1.36.0 and Swarm version 3.1.6 were run using default parameter settings. Because Swarm can produce small, low-abundance clusters, including singletons [30], we applied an additional post-processing step in which only non-singleton clusters with a centroid copy number of two or greater were retained.

### Assessment of clustering and denoising quality

For consistency, outputs from all clustering and denoising pipelines are referred to as clusters, including amplicon sequence variants (ASVs) inferred by DADA2 and denoised units inferred by UNOISE. Pipeline performance was evaluated using ground-truth-based metrics, including the total number of clusters, the number of perfect clusters, average cluster purity, the number of unique species represented in the output, the number of species represented by multiple clusters, concordance between estimated and true relative abundance profiles, and the number of reads not assigned to any output cluster.

A perfect cluster was defined as a cluster that was both 100% pure and 100% complete with respect to a single species. Thus, all reads within the cluster originated from that species, and all input reads originating from that species were assigned to the cluster. Here, “input reads” refers to the reads provided to the clustering or denoising step. Reads removed during preprocessing were excluded from this definition.

Average cluster purity was calculated as the average purity across all clusters, with cluster purity defined as the proportion of reads within a cluster belonging to the most abundant ground-truth species. To quantify the number of unique species represented in the output, each cluster was assigned a species label by majority voting based on the ground-truth taxonomy of its reads. Specifically, each cluster was assigned to the species corresponding to the most frequent ground-truth species label among the reads it contained. Using the same assignment procedure, we then counted the number of species represented by multiple clusters, defined as species assigned as the majority label in more than one cluster.

Concordance with the ground-truth community composition was assessed by calculating the Pearson correlation between the estimated and true relative abundance profiles. The estimated relative abundance of each species was defined as the total number of reads across clusters assigned to that species by majority voting, divided by the total number of clustered reads. Species present in the ground-truth community but not represented in any output cluster were assigned an estimated relative abundance of zero. The same evaluation framework was subsequently used to investigate how variation in the minimum abundance threshold affected the performance of the VSEARCH-based pipelines.

### Evaluating the effect of the minimum abundance threshold

The minimum abundance threshold (*minsize*) is a key parameter in VSEARCH-based pipelines because it determines which dereplicated sequences are retained for the primary clustering or denoising step. Because this threshold directly affects the balance between removing low-abundance sequences representing potential technical noise and retaining rare biological variants, its effect may be particularly important in highly diverse microbial communities. We therefore systematically varied *minsize* to evaluate how minimum abundance filtering in clustering and denoising affects the performance of VSEARCH-based pipelines across datasets of varying complexity. No directly equivalent minimum abundance parameter was evaluated for DADA2 in the pipelines used here.

Each simulated replicate dataset was analysed separately. Thus, dereplication and thresholding were performed within each dataset rather than after pooling reads across replicate datasets. This design allowed the effect of the minimum abundance threshold to be evaluated in relation to sequencing depth, as the copy number of the dereplicated sequences depends on the number of reads available within each dataset.

For both UNOISE and *cluster_size*, five minimum abundance thresholds were evaluated: 1, 2, 5, 10, and 20. For UNOISE, a *minsize* of 8 was additionally included because this represents the default setting for UNOISE. In all cases, the threshold refers to the minimum copy number required for a dereplicated sequence to be included directly in the primary clustering or denoising step. In the *cluster_size* pipeline, sequences below this threshold were subsequently mapped to the inferred centroids, as described above.

Because the effect of a fixed minimum abundance threshold may vary with sequencing depth, threshold analyses were performed at sequencing depths of 10 000, 50 000, and 100 000 reads. These analyses assessed how the minimum abundance threshold affected clustering and denoising performance, and whether its effects depended on sequencing depth. Performance was evaluated using the same metrics described above, and average values across replicate datasets were calculated for each configuration.

### Evaluation on real environmental data

To examine whether threshold-dependent patterns observed in simulated data were also reflected in empirical environmental data, we analysed 16S rRNA amplicon sequencing data from seafloor sediment samples from the AQUAeD project (https://www.nmbu.no/forskning/prosjekter/aquaed). The full dataset consisted of 300 samples from 50 sampling sites, with six replicate samples per site. For the present analysis, we used samples from ten randomly selected sites. Samples were sequenced using Illumina paired-end short-read technology targeting the 16S rRNA V3-V4 region. The sampling, DNA extraction, and sequencing protocols are described in detail by Philip [31]. Reads were preprocessed as described for the simulated datasets.

Because the true community composition was unknown, performance could not be evaluated using ground-truth-based metrics. Instead, repeated detection across replicate samples within each sampling site was used as an empirical indication of consistent biological signal. We therefore evaluated the effect of the minimum abundance threshold by comparing sequence prevalence profiles before and after clustering or denoising within each sampling site.

Prevalence was defined as the number of replicate samples within a sampling site in which a given dereplicated sequence was detected. To enable fair comparisons within sampling sites, replicate samples were rarefied to a common sequencing depth before analysis using a fixed random seed. Two rarefaction depths were used: 10 000 reads and the minimum sequencing depth observed among the replicate samples within each sampling site.

For UNOISE, thresholds of 1, 2, 5, 8, 10, and 20 were evaluated, whereas thresholds of 1, 2, 5, 10, and 20 were evaluated for *cluster_size*. Denoising was performed using *vs_cluster_unoise*, and the UC-format output was joined with the original input sequences to identify sequences that were assigned to clusters and those that remained unassigned. Sequences assigned to a cluster were considered retained, whereas sequences not assigned to any cluster were considered lost. For each sample, the original abundances of all sequences assigned to the same denoised cluster were summed. Cluster prevalence was then calculated as the number of replicate samples in which the summed cluster abundance was greater than zero. Sequences not assigned to any cluster were assigned zero abundance and zero prevalence.

Clustering was performed with sequences above the minimum abundance threshold using *vs_cluster_size*. The resulting centroids were used as references for global alignment of all reads using *vs_usearch_global*. Following global alignment, sequences assigned to the same centroid were represented by the corresponding cluster abundance within each sample, as returned by *vs_usearch_global*. Cluster prevalence was then calculated as the number of replicate samples in which this abundance was greater than zero. Sequences not assigned to any cluster were assigned zero abundance and zero prevalence.

The effect of the minimum abundance threshold was assessed by quantifying sequence loss. A sequence was considered lost if it was not assigned to any cluster and therefore had zero prevalence after processing.

## Results

### Clustering and denoising performance across simulated communities

The simulated datasets represented six community configurations comprising three levels of species richness (100, 1 000, and 10 000 species) and two maximum-to-minimum abundance ratios (10 and 100), with 20 replicate datasets generated for each configuration. Unless otherwise stated, results are based on datasets with a sequencing depth of 100 000 reads. Preprocessing of simulated reads showed consistently high read retention across datasets, with approximately 97% of reads retained after trimming and quality filtering, and 93-96% remaining after chimera removal (Supplementary Table S1).

We first compared pipeline performance based on the total number of clusters, pure clusters, perfect clusters, and ground-truth species retained after trimming and quality filtering (Figure 2). Differences among pipelines became more pronounced with increasing species richness, particularly in the total number of clusters produced and the number of perfect clusters.

**Figure 2.**
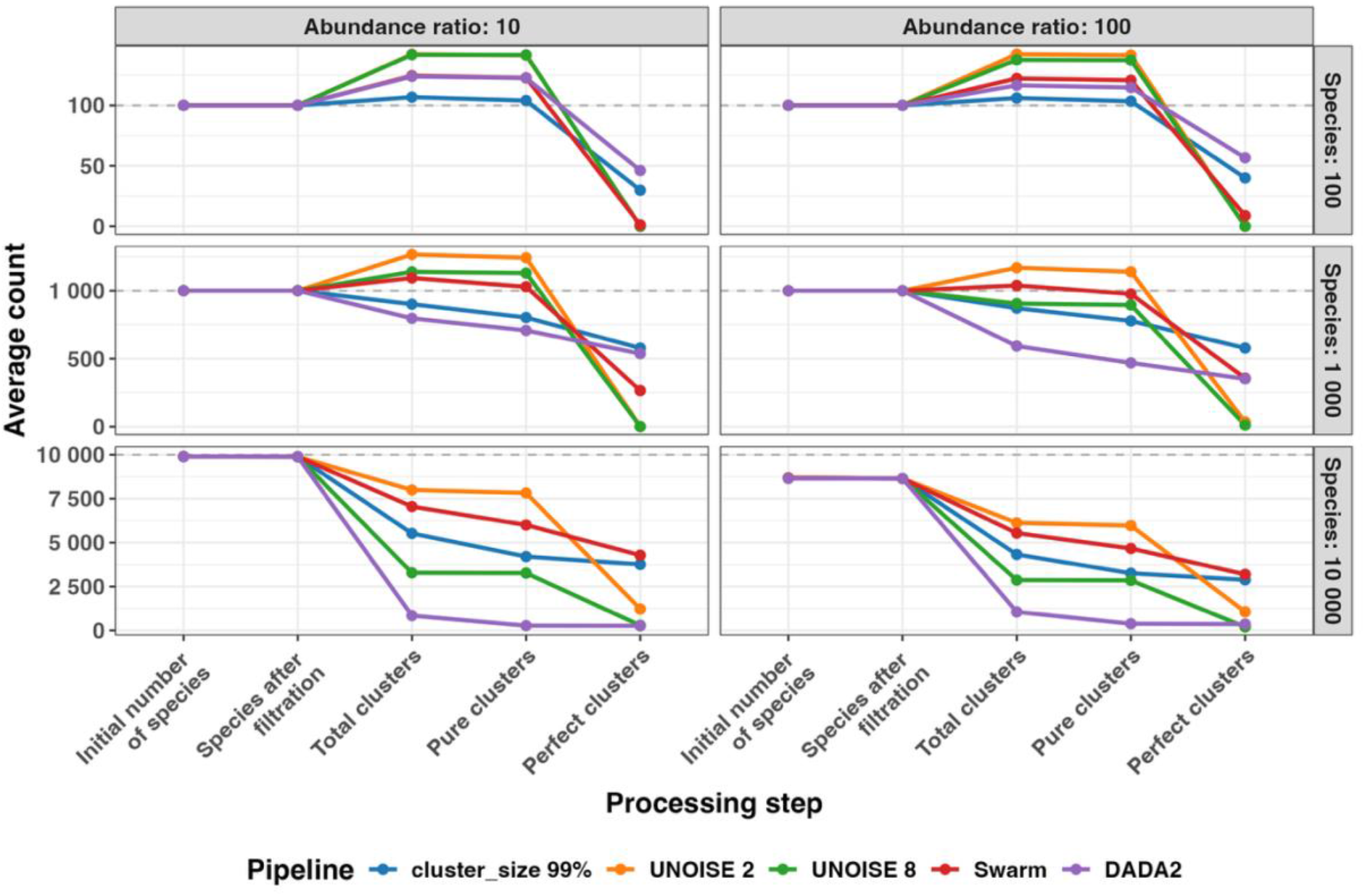
Average numbers of retained ground-truth species, total clusters, pure clusters, and perfect clusters across clustering and denoising pipelines at increasing levels of species richness and abundance ratio. Pure clusters contained sequences from a single species, whereas perfect clusters contained all and only the sequences originating from a single species. Points and lines represent averages across 20 replicate datasets for each combination of species richness and abundance ratio. The dotted line in each panel indicates the specified number of species in the community configuration.

At a species richness of 100, the average number of clusters ranged from 106 for *cluster_size* to 142 for UNOISE 2, compared with 100 simulated species. Most clusters were pure, whereas the number of perfect clusters was substantially lower. DADA2 produced the highest average number of perfect clusters, with 46 at an abundance ratio of 10 and 56 at an abundance ratio of 100, whereas the UNOISE configurations produced no perfect clusters.

At a species richness of 1 000, the average number of clusters ranged from 796 for DADA2 to 1 266 for UNOISE 2 at an abundance ratio of 10, and from 592 to 1 166 at an abundance ratio of 100. The number of pure clusters remained high but was slightly lower than in the 100-species datasets. At an abundance ratio of 10, DADA2 and *cluster_size* produced fewer clusters than the initial 1 000 species but generated the highest numbers of perfect clusters. At an abundance ratio of 100, *cluster_size* produced the highest average number of perfect clusters (578), whereas UNOISE 2 and UNOISE 8 produced none. Under this abundance ratio, only Swarm and UNOISE 2 produced more clusters than the initial species count.

At a species richness of 10 000, all pipelines produced fewer clusters than the initial number of species. At an abundance ratio of 10, average cluster counts ranged from 844 for DADA2 to 7 996 for UNOISE 2, whereas at an abundance ratio of 100 they ranged from 1 056 to 6 126. The UNOISE configurations showed a close correspondence between total and pure cluster counts but generated substantially fewer perfect clusters than *cluster_*size and Swarm. Counts were generally lower at an abundance ratio of 100 for most pipelines. Under this condition, the number of ground-truth species represented after preprocessing was also lower than the specified species richness, because some low-abundance species received no reads during simulation.

Overall, deviation from the ideal one-species-one-cluster pattern increased with species richness.

### Effects of species richness and abundance ratio on average cluster purity

We next assessed whether the observed differences in cluster numbers were accompanied by differences in average cluster purity. Average cluster purity remained high across datasets, although differences among pipelines became more pronounced with increasing species richness (Figure 3). Within each panel, variation among replicate datasets was limited for each pipeline, as indicated by the narrow boxplots. At a species richness of 100, average cluster purity ranged from 0.98 to 1.0 across pipelines and abundance ratios. At a species richness of 1 000, purity remained high, ranging from 0.95 to 0.99 across pipelines, with UNOISE 2 and UNOISE 8 showing the highest values. At a species richness of 10 000, both UNOISE configurations maintained high average purity, with values of approximately 0.99, whereas *cluster_size*, Swarm, and particularly DADA2 showed lower values. Across abundance ratios, average cluster purity at this richness level ranged from 0.63 for DADA2 to 0.99 for UNOISE 2 and UNOISE 8. Overall, average cluster purity decreased with increasing species richness for all pipelines except UNOISE 2 and UNOISE 8, although the magnitude of the decline varied among pipelines. Because high average cluster purity may coexist with species splitting across multiple clusters, we next examined species representation and cluster splitting directly.

**Figure 3.**
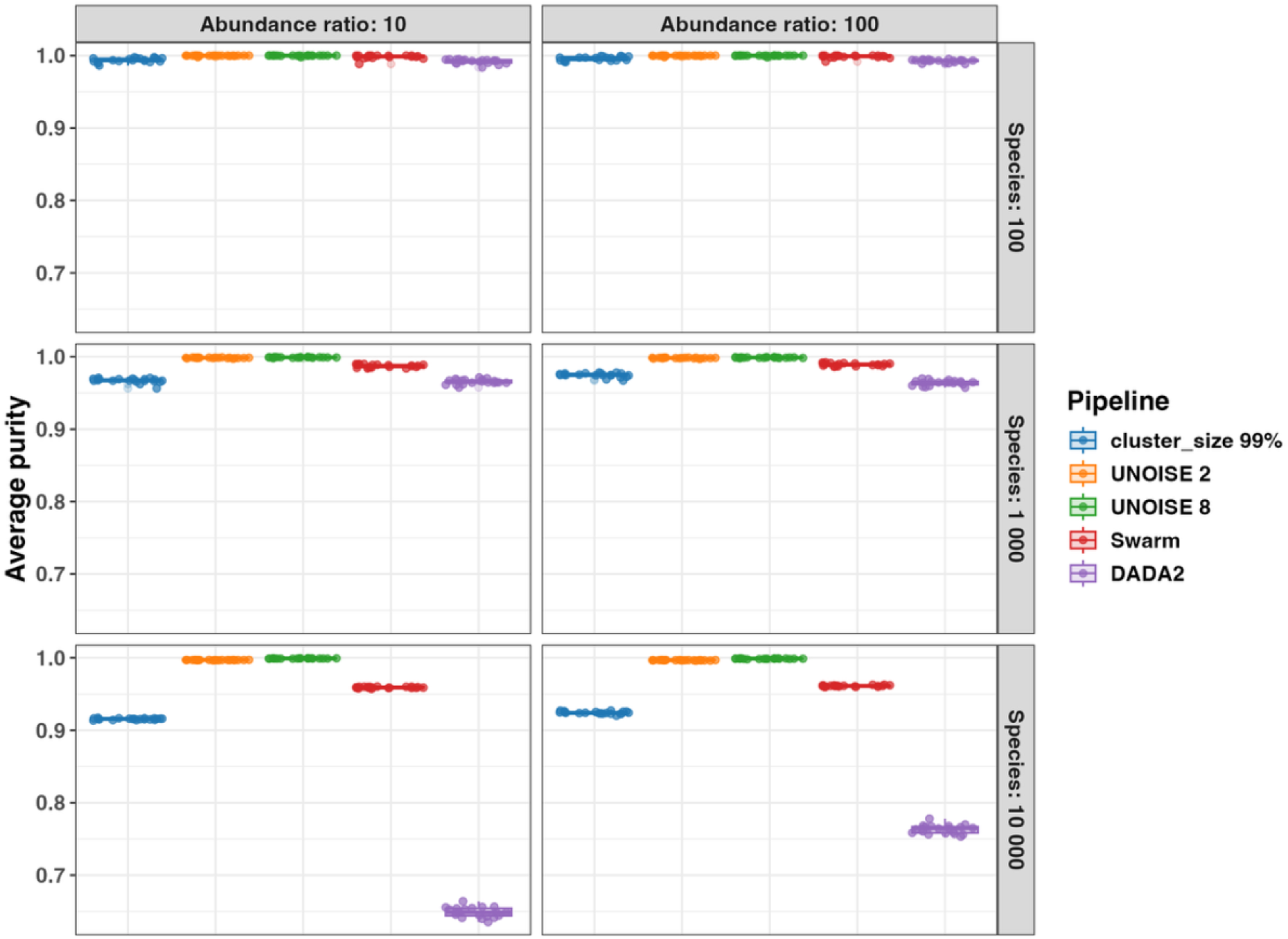
Average cluster purity across clustering and denoising pipelines at increasing levels of species richness and abundance ratio. Boxplots summarise results across 20 replicate datasets for each combination of species richness and abundance ratio. A cluster purity of 1.0 represents the optimal value and indicates that all sequences within a cluster originate from the same species.

### Species representation and splitting across clusters

We next examined the balance between species representation and splitting across multiple clusters (Figure 4). The ideal outcome was high species representation combined with limited splitting, corresponding to the point where the two dotted lines meet near the lower-right corner of each panel.

**Figure 4.**
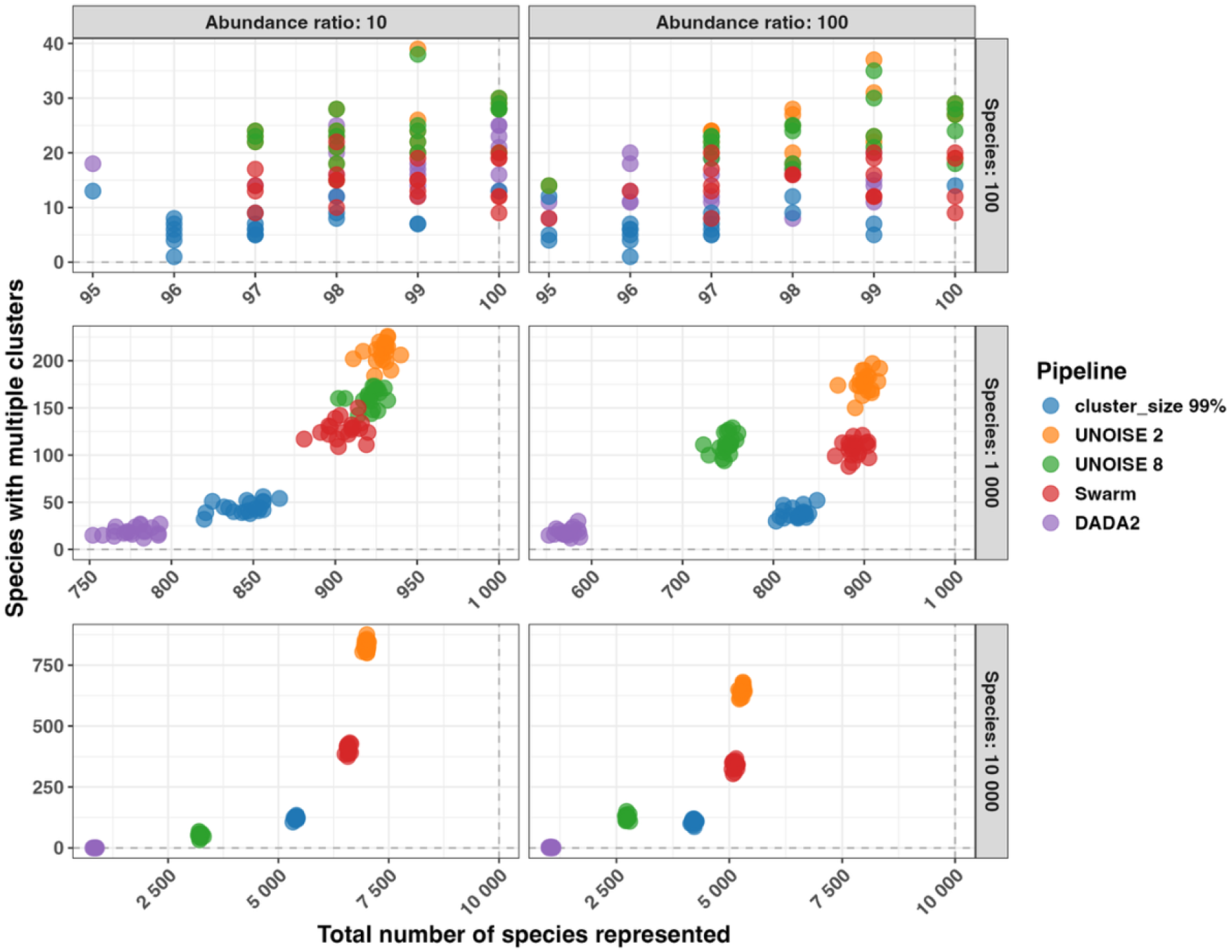
Species representation and splitting across clusters for clustering and denoising pipelines at increasing levels of species richness and abundance ratio. Species representation was defined as the number of ground-truth species assigned as the majority label of at least one cluster. Species splitting was defined as the number of represented species assigned as the majority label of more than one cluster. Points represent individual replicate datasets. The vertical dotted line indicates the specified number of species in the simulated community, and the horizontal dotted line indicates zero splitting.

At a species richness of 100, all pipelines represented 95-100 species but differed in the extent of splitting. The average number of species split across multiple clusters ranged from 1 for *cluster_size* to 39 for UNOISE 2, with Swarm and DADA2 showing intermediate values. No pipeline consistently represented all 100 species across replicates.

At a species richness of 1 000, differences among the pipelines became more pronounced and varied with abundance ratio. At an abundance ratio of 10, average species representation ranged from 752 for DADA2 to 940 for UNOISE 2, while an average of 19 and 209 species, respectively, were split across multiple clusters. At an abundance ratio of 100, representation decreased to 553-917 across pipelines. DADA2 again represented the fewest species and produced the least splitting, whereas UNOISE 2 represented the most species and produced the greatest splitting. Replicate datasets formed tight pipeline-specific groups, indicating limited within-pipeline variation.

At a species richness of 10 000, the trade-off between species representation and splitting became more pronounced. At an abundance ratio of 10, average species representation ranged from 845 for DADA2 to 6 990 for UNOISE 2, compared with 1 055 to 5 280 species at an abundance ratio of 100. The separation between UNOISE 2 and UNOISE 8 was particularly pronounced at this richness level, with UNOISE 2 representing more species but also splitting substantially more species across clusters. Swarm represented nearly as many species as UNOISE 2, but with less splitting. *cluster_size* represented more species than UNOISE 8 while also splitting fewer species across both abundance ratios. None of the pipelines represented all 10 000 species, and species representation was generally lower in the more uneven communities.

Together, these results indicate a trade-off between species representation and species splitting, with the magnitude of this trade-off depending on both species richness and abundance ratio. We next evaluated reconstruction of the underlying abundance structure of the simulated datasets.

### Reconstruction of ground-truth abundance profiles

We then evaluated how well each pipeline reconstructed the ground-truth abundance profiles of the simulated communities (Figure 5). At a species richness of 100, all pipelines showed strong concordance with the ground-truth abundance profiles, with correlations ranging from 0.87 to 1.0 across pipelines. At a species richness of 1 000, differences among pipelines became more pronounced, with average correlations ranging from approximately 0.52 for *cluster_size* to 0.97 for DADA2 across abundance ratios. At a species richness of 10 000, correlations decreased markedly, ranging from approximately 0.25 for *cluster_size* to 0.55 for the UNOISE configurations. UNOISE 2, UNOISE 8, and DADA2 showed the highest correlations at this richness level, whereas *cluster_size* showed substantially lower concordance with the ground-truth abundance profiles. Overall, abundance ratio had relatively little effect at low species richness, whereas differences between abundance ratios became more apparent at higher richness, with generally higher correlations at an abundance ratio of 100.

**Figure 5.**
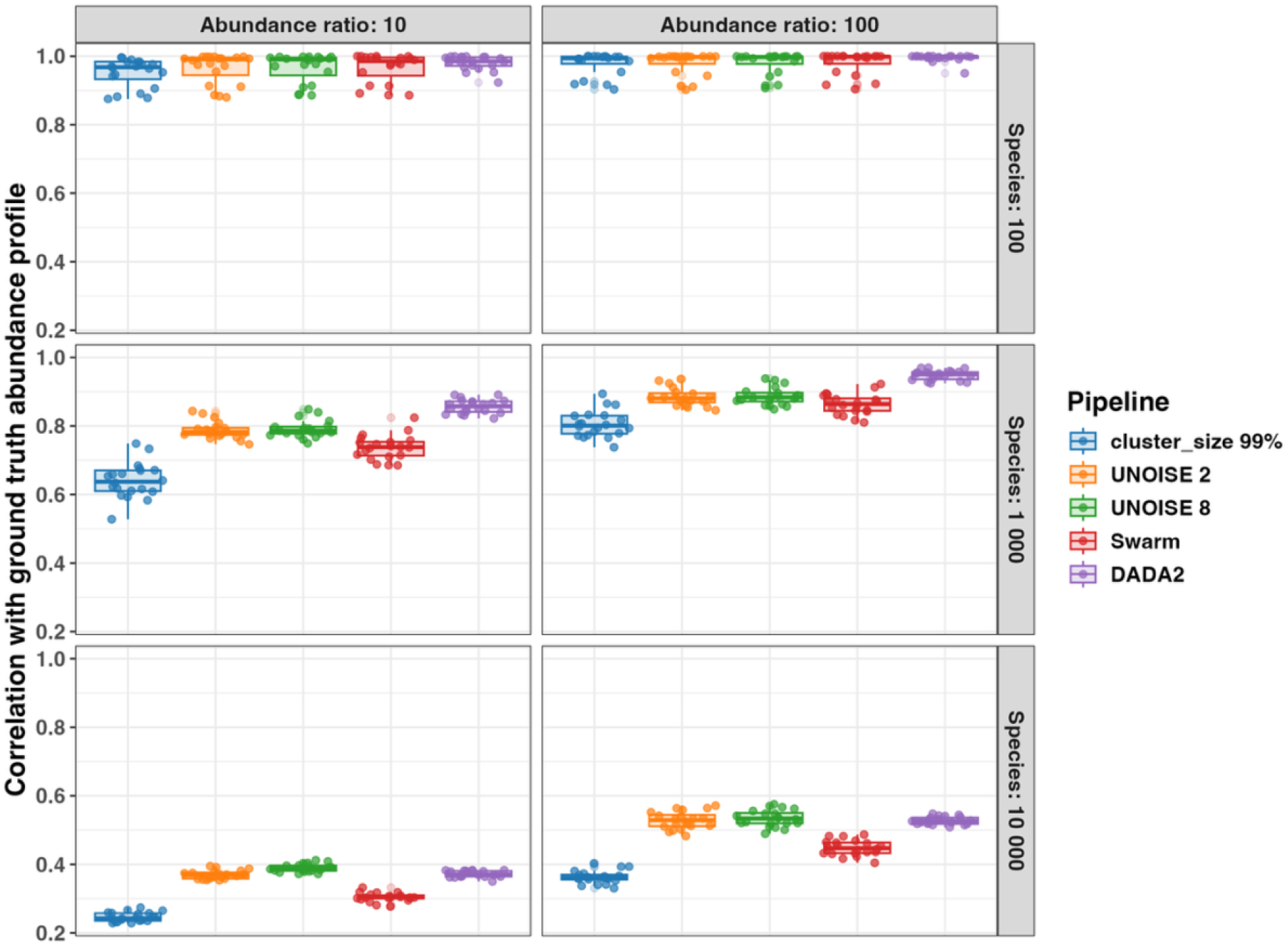
Correlation between estimated and ground-truth relative abundance profiles across clustering and denoising pipelines at increasing levels of species richness and abundance ratio. Boxplots summarise Pearson correlation coefficients across 20 replicate datasets for each combination of species richness and abundance ratio. A correlation of 1.0 indicates perfect concordance with the ground-truth relative abundance profile.

### Unclustered reads across pipelines

Because reads could remain unassigned through pipeline-specific filtering, denoising, remapping, or post-processing steps, we examined the number and ground-truth species origin of unclustered reads. The number of unclustered reads varied substantially among pipelines across all combinations of species richness and abundance ratio (Figure 6), whereas variation among replicate datasets was generally limited. At a species richness of 100, the average number of unclustered reads ranged from 163 for *cluster_size* to 20 850 for UNOISE 8. At a species richness of 1 000, average values ranged from 527 to 21 887 reads. At a species richness of 10 000, the pattern changed with DADA2 producing the highest average number of unclustered reads (more than 40 000), compared with approximately 2 000 to 3 000 reads for *cluster_size* and Swarm. Overall, the number of unclustered reads differed substantially among pipelines and varied with species richness, with the UNOISE configurations producing the highest numbers at lower richness levels and DADA2 at the highest richness level.

**Figure 6.**
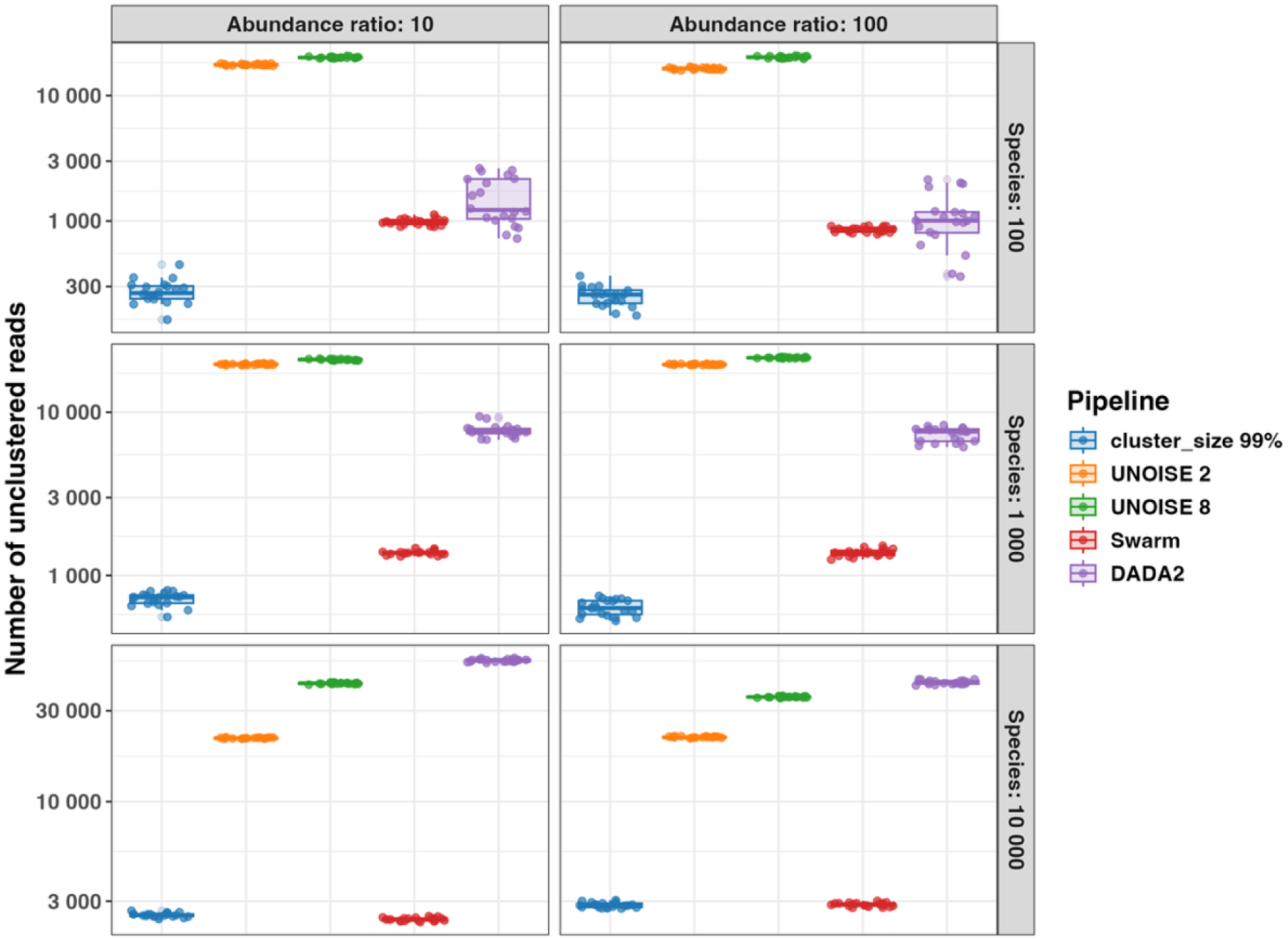
Number of unclustered reads across clustering and denoising pipelines at increasing levels of species richness and abundance ratio. Boxplots summarise results across 20 replicate datasets for each combination of species richness and abundance ratio. The y-axis scale varies among species-richness levels; absolute vertical distances should therefore not be compared across rows.

Species representation among unclustered reads also varied substantially among pipelines (Figure 7), whereas variation among replicate datasets was generally limited. Across all species-richness levels and abundance ratios, UNOISE 2 and UNOISE 8 had the highest numbers of species represented among unclustered reads. At a species richness of 100, the average number of represented species ranged from approximately 50-100 across pipelines, with *cluster_size* and DADA2 generally showing the lowest values.

**Figure 7.**
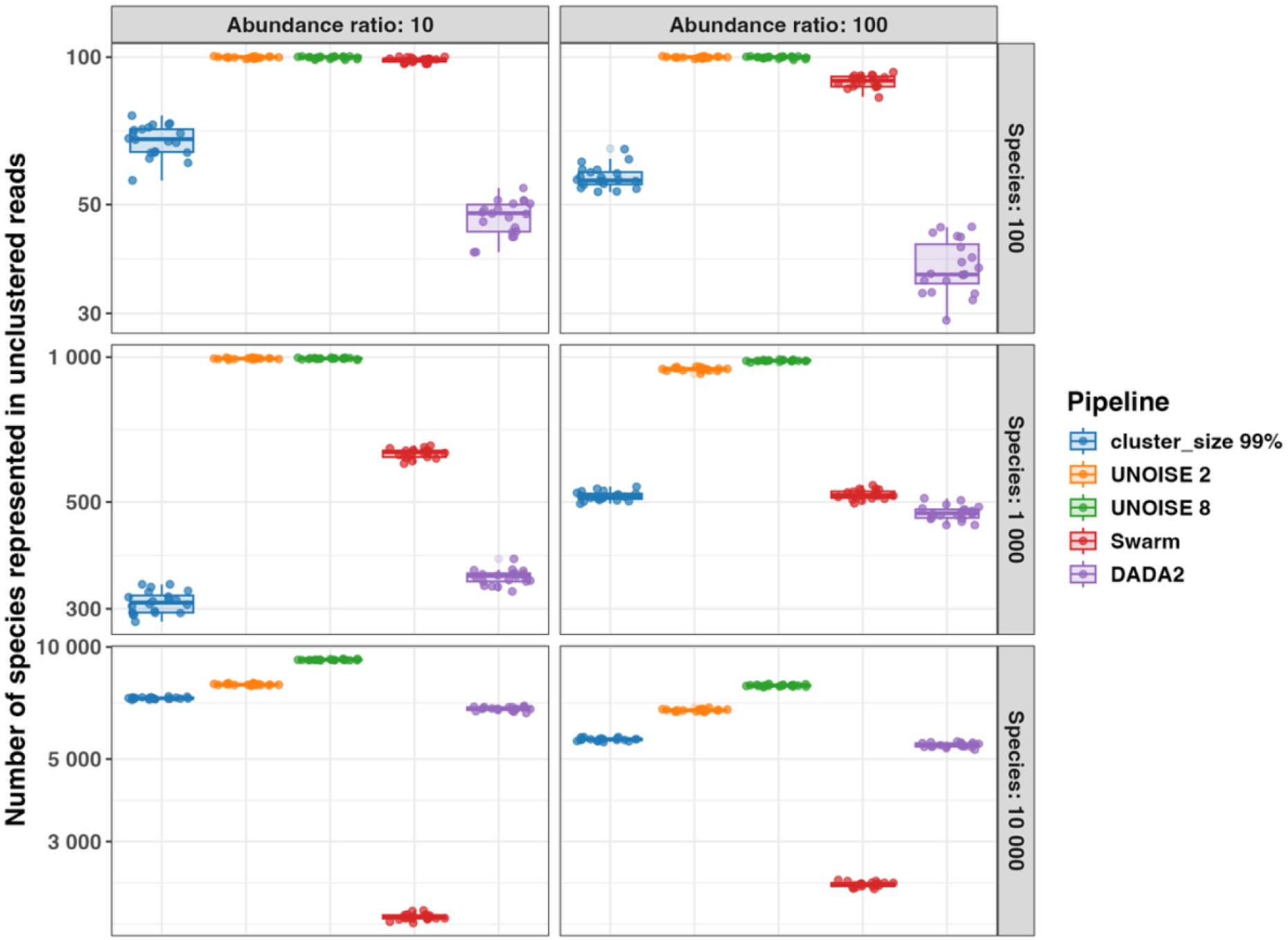
Number of species represented among unclustered reads across clustering and denoising pipelines at increasing levels of species richness and abundance ratio. The figure shows the number of ground-truth species represented among reads not assigned to any cluster. Boxplots summarise results across 20 replicate datasets for each combination of species richness and abundance ratio. The y-axis scale varies among species-richness levels; absolute vertical distances should therefore not be compared across rows.

At a species richness of 10 000, differences among pipelines were more pronounced. Swarm had the fewest species represented among unclustered reads, with average values of approximately 1 800 and 2 200 at abundance ratios of 10 and 100, respectively, compared with 5 000 to 10 000 for the other pipelines. At this richness level, UNOISE 8 had more represented species than UNOISE 2 at both abundance ratios.

Overall, unclustered reads originated from a broad range of ground-truth species for most pipelines.

We further examined the ground-truth abundances of species represented among unclustered reads. Ground-truth abundance distributions indicated that unclustered reads generally originated from low-abundance species (Supplementary Figure S1).

### Evaluating the effect of the minimum abundance threshold

Given the observed pipeline-specific differences in species representation and cluster formation, we next examined how the minimum abundance threshold affects VSEARCH-based pipelines at sequencing depths of 10 000, 50 000, and 100 000 reads. Figure 8 summarises four representative metrics for the dataset comprising 10 000 species and an abundance ratio of 10: the number of clusters, the number of perfect clusters, the number of species split across multiple clusters, and the correlation with the ground-truth relative abundance profile.

**Figure 8.**
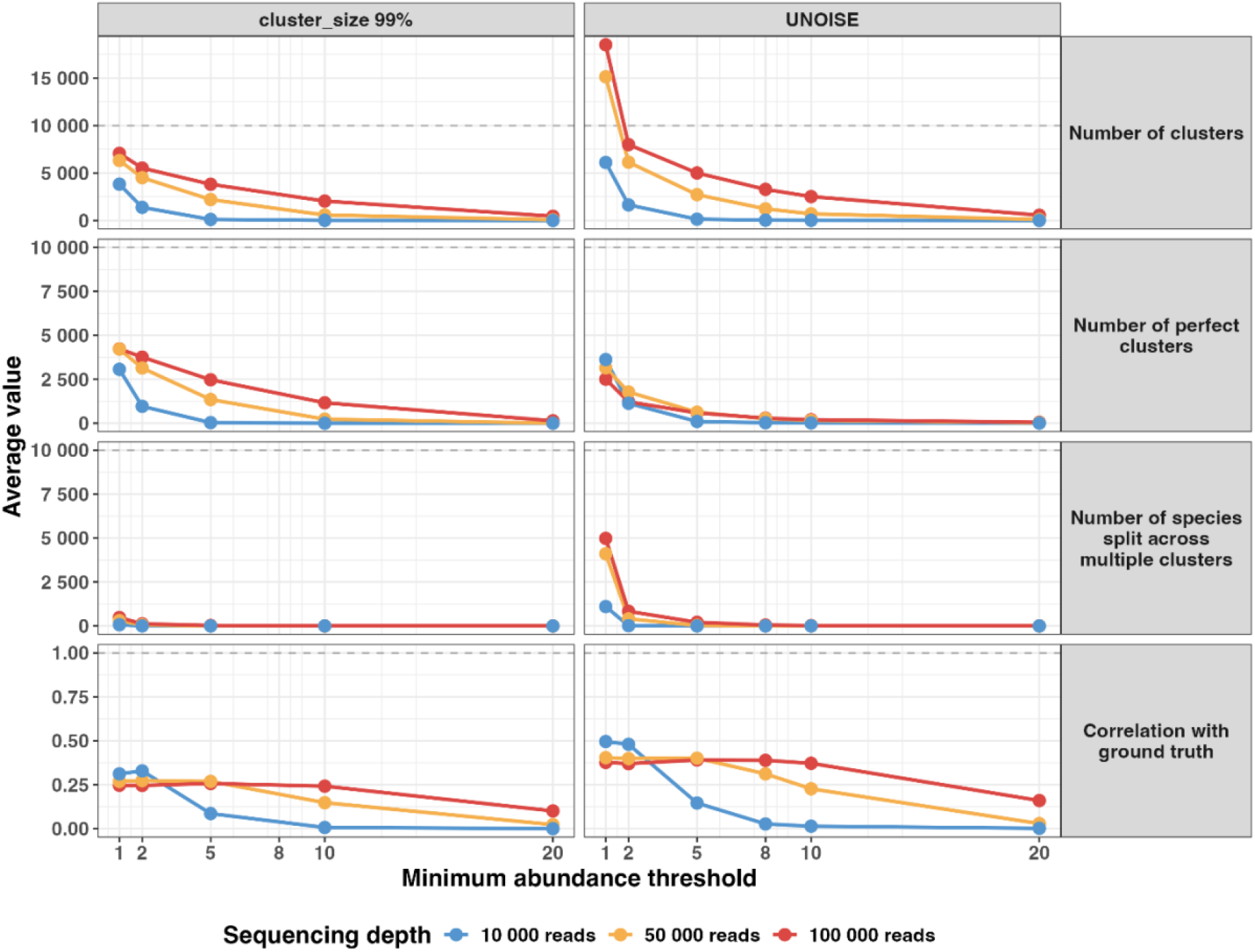
Effects of minimum abundance threshold and sequencing depth on the performance of VSEARCH-based clustering and denoising pipelines for the dataset containing 10 000 species and an abundance ratio of 10. Panels compare cluster_size at 99% sequence identity and UNOISE across minimum abundance thresholds and sequencing depths of 10 000, 50 000, and 100 000 reads. Points and lines represent average values across 20 replicate datasets for each configuration. The horizontal dashed lines indicate the ground-truth species count or the optimal value, where applicable.

For both *cluster_size* and UNOISE, increasing the minimum abundance threshold reduced the number of clusters. At a threshold of 1, UNOISE produced approximately 15 000 and 18 500 clusters at sequencing depths of 50 000 and 100 000 reads, respectively, exceeding the 10 000 simulated species under both conditions. In contrast, *cluster_size* produced fewer than 10 000 clusters at all evaluated thresholds and sequencing depths, ranging from approximately 3 800 to 7 000 clusters at threshold 1. The number of perfect clusters also declined with increasing threshold, approaching zero for UNOISE at higher thresholds despite the low number of species split across multiple clusters. The number of species split across multiple clusters decreased most strongly between thresholds 1 and 2. For example, at 100 000 reads, UNOISE decreased from approximately 5 000 species split at threshold 1 to approximately 800 at threshold 2, compared with approximately 480 and 120 for *cluster_size*.

Correlation with the ground-truth relative abundance profile also decreased with increasing minimum abundance threshold, although the magnitude of this effect depended on sequencing depth. At a sequencing depth of 10 000 reads, correlations declined at relatively low thresholds. At 50 000 reads, correlations remained relatively stable across a wider range of thresholds, whereas at 100 000 reads, substantial declines occurred only at higher thresholds. Overall, increasing the minimum abundance threshold had a greater effect on reconstruction of relative abundance profiles at lower sequencing depths. At the highest thresholds, the number of clusters, number of perfect clusters, number of species split across multiple clusters, and correlation with the ground-truth abundance profile all declined markedly, with some values approaching zero. The remaining datasets, representing other combinations of species richness and abundance ratio, showed broadly similar trends (Supplementary Figures S2-S6), while results for the remaining metrics are provided in Supplementary Table S2.

### Evaluation using real environmental data

To examine whether the threshold-dependent patterns observed in the simulated datasets were also reflected in empirical environmental data, we applied the same range of minimum abundance thresholds to 16S rRNA amplicon data from seafloor sediment samples (Figure 9). Because these data lacked ground truth, we did not evaluate accuracy directly. Instead, we examined threshold-dependent sequence retention across prevalence levels within sampling sites. Prevalence was defined as the number of replicate samples in which a sequence was detected, and retention as the percentage of unique input sequences within each original prevalence level that remained assigned after processing. Sequences detected in one to three replicates were classified as low prevalence, whereas those detected in four to six replicates were classified as high prevalence.

**Figure 9.**
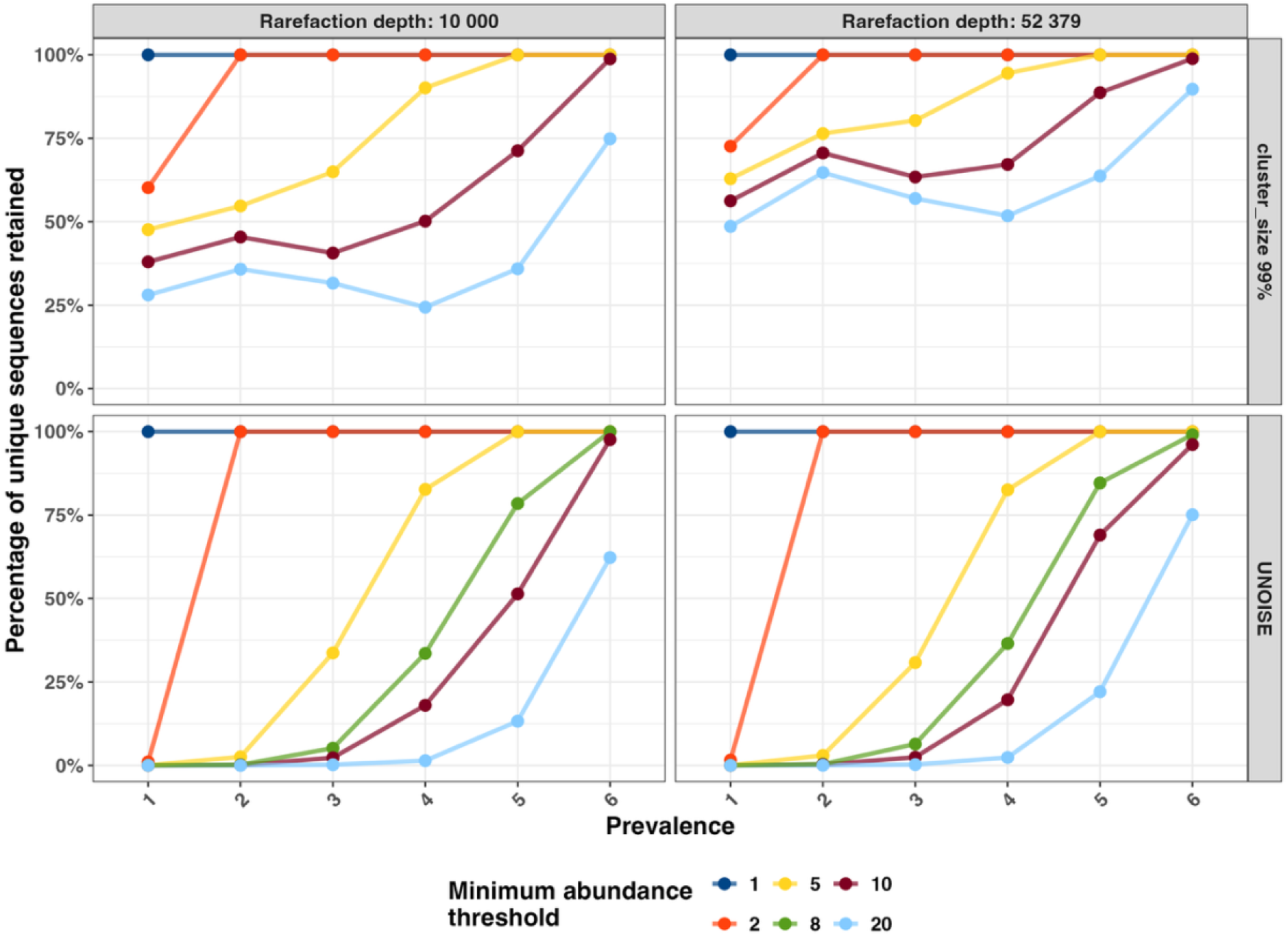
Retention of unique input sequences across prevalence levels following the application of different minimum abundance thresholds to seafloor sediment data. The figure shows the percentage of unique input sequences retained after clustering with VSEARCH *cluster_size* or denoising with UNOISE within one sampling site. Prevalence was defined as the number of replicate samples within the sampling site in which a sequence was detected before clustering or denoising. Results are shown at rarefaction depths of 10 000 reads and 52 379 reads, the latter corresponding to the minimum sequencing depth among replicate samples within the sampling site.

Increasing the minimum abundance threshold reduced sequence retention in both *cluster_size* and UNOISE, although the magnitude of this effect differed between pipelines. At a threshold of 1, both pipelines retained approximately 100% of sequences across all prevalence levels. At a threshold of 2, the pipelines differed markedly among sequences detected in only one replicate: UNOISE retained almost none of these sequences, whereas *cluster_size* retained a substantial proportion.

Retention generally increased with prevalence at thresholds of 5 or higher, particularly for UNOISE. However, sequence loss also occurred among high-prevalence sequences. Increasing the threshold reduced the retention of sequences detected in four to six replicate samples, and even sequences detected in all six replicates could be excluded at the strictest threshold. This effect was most pronounced at a rarefaction depth of 10 000 reads.

Th**e** difference in retention between the two rarefaction depths was greatest at prevalence levels of 3**-**5 and at stricter minimum abundance thresholds. Greater rarefaction depth reduced, but did not eliminate, the loss of high-prevalence sequences. Compared with *cluster_size*, UNOISE generally retained fewer sequences at thresholds of 5 or higher, particularly at lower prevalence levels. These empirical results were consistent with the simulated-data analyses, showing that minimum abundance thresholds can remove repeatedly detected sequences and that this effect depends on sequencing depth, prevalence, and pipeline. Results from the remaining sampling sites showed broadly similar trends and are provided in Supplementary Table S3.

## Discussion

In this study, we benchmarked four clustering and denoising pipelines for 16S rRNA amplicon data using simulated microbial communities with known ground-truth compositions. By varying species richness, abundance ratio, sequencing depth, and minimum abundance thresholds, we assessed how pipeline performance changed across dataset conditions. Overall, performance varied with species richness, abundance ratio, evaluation metric, and parameter selection. Differences among pipelines were modest in communities with low species richness but became substantially greater as species richness increased. These findings suggest that pipeline and parameter selection may be considerably more important for highly diverse environmental datasets. Previous comparisons of environmental 16S rRNA workflows have likewise shown that pipeline choice can affect recovered diversity and the ecological interpretation [13], [14]. This underscores the need for careful consideration and transparent reporting of both pipeline selection and parameter settings when analysing complex environmental communities.

No pipeline consistently combined high cluster purity, species representation, perfect-cluster recovery, and strong concordance with relative abundance profiles with limited species splitting and read loss across all dataset conditions, indicating clear trade-offs in pipeline selection for high-diversity environmental samples. These trade-offs were relatively minor in the 100-species datasets, where the pipelines produced broadly similar results across several metrics, but became increasingly pronounced at species-richness levels of 1 000 and 10 000. Although abundance ratio affected several metrics, differences in pipeline performance were more consistently pronounced across levels of species richness. This supports the premise that performance observed in low-complexity or mock-community datasets may not reliably predict pipeline behaviour in highly diverse environmental samples. Recent benchmarking using mock communities has similarly demonstrated method-specific differences in over-splitting and over-merging [16]. Our results extend these observations to substantially higher species richness, where differences among pipelines became increasingly pronounced. In particular, the increasing deviation from the ideal one-species-one-cluster pattern with increasing species richness suggests that highly diverse communities amplify differences in how pipelines handle closely related sequences, low-abundance variants, and sequencing errors. However, the one-species-one-cluster pattern should be considered an analytical reference rather than an expected biological outcome, because species boundaries cannot always be resolved using 16S rRNA amplicon sequences alone [16]. In addition, individual bacterial genomes may contain multiple, slightly different copies of the 16S rRNA gene, which can cause a single species to be represented by more than one sequence variant or cluster [32].

A central finding was that high average cluster purity did not necessarily imply accurate species-level reconstruction. Although several pipelines produced clusters with high average purity, indicating that most clusters were dominated by reads from a single ground-truth species, the number of perfect clusters was consistently much lower. This suggests that many species were split across multiple high-purity clusters, were incompletely represented after preprocessing and clustering, or both. The implications of this trade-off depend on the intended downstream application. Species splitting may have limited consequences when the resulting clusters are ultimately assigned to the same taxonomic group, but it becomes more problematic for analyses requiring accurate estimates of richness, abundance, or community composition. The UNOISE-based pipelines illustrate this trade-off particularly clearly: they maintained high average cluster purity but produced very few perfect clusters. High purity alone may therefore provide an overly favourable impression of pipeline performance if completeness and species splitting are not considered simultaneously.

Rather than identifying a single pipeline that performed best across all metrics, our results revealed complementary performance profiles reflecting different analytical priorities.

UNOISE, particularly UNOISE 2, may be favoured when high species representation and cluster purity are primary objectives, while also providing comparatively strong concordance with ground-truth relative abundance profiles, especially in highly diverse datasets. However, this comes at the cost of increased species splitting and a substantial number of unclustered reads, which may be disadvantageous when closer correspondence between biological species and individual output features is desired. One contributing factor is that reads below the abundance threshold were not globally remapped to the denoised output features and were therefore excluded from the final output.

DADA2 may be useful when limited species splitting is prioritised, while also providing strong reconstruction of relative abundance profiles comparable to UNOISE. However, these characteristics were accompanied by lower species representation at high richness and lower average cluster purity in several configurations. Thus, despite strong performance for some metrics, DADA2 was not consistently the optimal choice in highly diverse communities.

VSEARCH *cluster_size* also limited species splitting and may therefore be useful when reducing species splitting is a primary objective. However, its generally weaker reconstruction of ground-truth abundance profiles suggests that this advantage should be weighed against reduced performance for abundance-based analyses.

Swarm showed a comparatively balanced performance profile in several high-richness datasets. It retained relatively high species representation while producing substantially less splitting than UNOISE 2, without showing the pronounced loss of species representation observed for DADA2. Swarm may therefore represent a useful compromise when both species representation and limited splitting are important, although it was not consistently the strongest pipeline for reconstruction of relative abundance profiles.

Previous benchmarks using lower-richness communities have likewise reported pipeline-specific differences in sensitivity and specificity, with DADA2 showing high sensitivity and UNOISE favouring greater specificity or a different balance between sensitivity and resolution [10], [12]. The different performance profiles observed at high species richness in the present study further indicate that relative pipeline performance depends on dataset characteristics and the metric being evaluated.

The number of unclustered reads varied substantially among pipelines and with species richness. Importantly, these reads were not restricted to a small subset of species but represented a broad range of ground-truth species and generally originated from low-abundance taxa. This pattern suggests that reads from rare members of the community are particularly vulnerable to exclusion during clustering and denoising. This is especially relevant in highly diverse environmental datasets, where rare taxa may represent genuine members of the microbial rare biosphere [33], while low-abundance sequence variants may also include spurious features introduced during sequencing or processing [34].

Distinguishing genuine low-abundance taxa from technical artefacts therefore remains a challenge in 16S rRNA amplicon analysis. Unclustered reads could therefore reflect the loss of low-abundance sequence information during bioinformatic processing rather than biological absence.

The minimum abundance threshold introduced a clear trade-off between reducing species splitting and retaining species representation and abundance information. Increasing the threshold reduced the total number of clusters and the number of species split across multiple clusters, but also reduced the number of perfect clusters, and at higher thresholds, the concordance with the ground-truth relative abundance profile. Because perfect clusters were defined as both pure and complete, their decline at higher minimum abundance thresholds may reflect reduced cluster completeness, cluster purity, or both. Importantly, the effect of the minimum abundance threshold became less pronounced with increasing sequencing depth. At lower sequencing depths, more dereplicated sequences have low copy numbers and are therefore more likely to fall below a given threshold. The importance of sequencing depth for the recovery of low-abundance community members is consistent with previous environmental 16S rRNA studies showing that sequencing depth can substantially influence observed microbial richness [35]. The same threshold can consequently have substantially different effects depending on sequencing depth. These results indicate that minimum abundance thresholds should be considered in relation to sequencing depth rather than assumed to perform equivalently across datasets. More broadly, previous benchmarking work has similarly emphasised that bioinformatic parameter settings should be evaluated in relation to the specific dataset and study design rather than applied without validation [9].

The focus on minimum abundance thresholds in VSEARCH-based pipelines was further motivated by previous work showing strong performance of UNOISE and Swarm relative to DADA2 in high-diversity marine seafloor samples [14]. However, the present results show that strong performance under one set of evaluation criteria does not imply consistently optimal performance across all metrics. Minimum abundance thresholds were evaluated only for the VSEARCH-based pipelines, where they directly determine which dereplicated sequences enter the primary clustering or denoising step. No directly equivalent threshold was evaluated for DADA2 or Swarm. Although DADA2 includes parameters such as *MIN_ABUNDANCE*, this parameter is primarily intended to reduce computational demands in large pooled datasets rather than to adjust biological sensitivity [36].

The analysis of real seafloor sediment extended the simulation results by showing that threshold-dependent sequence loss also occurred in empirical environmental data. Because these data lacked ground truth, excluded sequences cannot be classified as either true biological variants or technical artefacts. Nevertheless, sequence loss was not restricted to isolated, low-prevalence observations. Sequences detected across multiple samples within the same sampling site were also excluded at stricter thresholds, including at the commonly used default UNOISE setting of *minsize* = 8. Previous work has similarly shown that abundance-based filtering can improve the reproducibility of low-abundance feature detection across replicate samples, but at the cost of removing sequence features and affecting richness-sensitive diversity estimates [37]. These findings highlight the inherent trade-off between removing potentially unreliable low-abundance features and retaining sequence variation detected across multiple replicate samples. Threshold effects should therefore be interpreted in relation to sequencing depth, community properties, and the objectives of the study.

To ensure a fair comparison of clustering and denoising performance, all pipelines were provided with reads subjected to the same initial preprocessing. This reduced potential confounding from pipeline-specific trimming and quality filtering and allowed downstream differences to be attributed more directly to the clustering and denoising procedures. However, this standardisation also meant that pipelines were not evaluated as complete default workflows. This limitation was particularly relevant to DADA2, which includes its own trimming and filtering functions.

Several limitations should be considered when interpreting these results. First, although simulations provide a reproducible framework for assessing pipeline performance when empirical ground truth is unavailable, the simulated communities used here were simplified approximations of real environmental microbial communities. A strength of the simulation design was that all available 16S rRNA variants from the selected genomes were retained, thereby incorporating both within-species and within-genome sequence variation. This allowed the simulations to capture a broader range of biologically occurring 16S rRNA variation than would be represented by selecting a single sequence variant per genome. Nevertheless, the simulations represented environmental community complexity primarily through species richness and abundance ratio. Additional factors, including the limited species-level resolution of short 16S rRNA amplicon regions such as V3-V4, may further affect pipeline performance in empirical datasets [38]. Second, the one-species-one-cluster expectation used in the simulated benchmark should be regarded as an analytical reference point rather than a universally attainable biological outcome. Third, Pearson correlation measures concordance between relative abundance profiles but may not fully capture errors affecting rare taxa or species-level recovery. Finally, because the seafloor sediment analysis lacked ground truth, accuracy could not be evaluated directly. These data were therefore used only to assess whether the threshold-dependent retention patterns observed in simulations were also reflected in empirical data.

These findings have several methodological implications for high-diversity 16S rRNA amplicon studies. First, pipeline performance should not be evaluated based on a single metric. Average cluster purity, the number of perfect clusters, the number of represented ground-truth species, splitting across clusters, unclustered reads, and reconstruction of relative abundance profiles each capture distinct aspects of performance. Second, default parameter settings may not be appropriate for datasets that differ in sequencing depth or community composition. Stringent abundance thresholds can reduce apparent noise and species splitting but may also exclude reads from rare taxa. Third, pipeline selection should reflect the primary analytical objective. UNOISE may be favoured when species representation and cluster purity are prioritised, DADA2 when reconstruction of relative abundance profiles and limited splitting are more important, and Swarm when a compromise between species representation and splitting is desired. VSEARCH *cluster_size* may be useful when limited splitting is prioritised, although its weaker reconstruction of relative abundance profiles should be considered. These recommendations should be interpreted in the context of the simulated community structures and parameter settings evaluated here rather than as universal rankings of the pipelines.

## Conclusion

This study demonstrates that pipeline performance in highly diverse 16S rRNA amplicon datasets varies with species richness, abundance ratio, evaluation metric, and parameter selection. Simulated communities with known ground-truth composition showed that differences among pipelines became more pronounced as species richness increased, and that no single pipeline performed best across all metrics. High average cluster purity did not necessarily correspond to accurate species-level recovery, highlighting trade-offs among cluster purity, species representation, species splitting, relative abundance reconstruction, and read retention. Among the evaluated pipelines, UNOISE favoured high species representation and cluster purity while also showing strong reconstruction of relative abundance profiles, although with greater species splitting and read loss. DADA2 showed similarly strong reconstruction of relative abundance profiles with less splitting but lower species representation and cluster purity at high species richness. Swarm provided a balance between species representation and splitting, whereas VSEARCH *cluster_size* limited splitting but showed weaker reconstruction of relative abundance profiles. These distinct performance profiles indicate that pipeline selection should reflect the primary analytical objective. In the VSEARCH-based pipelines, the effects of the minimum abundance threshold varied with sequencing depth, while the seafloor sediment analysis showed that stricter thresholds could also remove sequences detected repeatedly across replicate samples. Together, these findings emphasise the importance of considering dataset characteristics and analytical objectives when selecting pipelines and parameter settings, and of transparently reporting analytical choices in studies of highly diverse environmental microbial communities.

## Declarations

### Ethics approval and consent to participate

Not applicable.

### Consent for publication

Not applicable.

### Availability of data and material

The real environmental sequencing data supporting the conclusions of this article are available in the European Nucleotide Archive (ENA) under the BioProject number PRJNA1128851 (https://www.ebi.ac.uk/ena/browser/view/PRJNA1128851). All scripts and associated files required to reproduce the simulated datasets and perform all analyses are available on GitHub (https://github.com/CassandraHjo/benchmarking-supplementary-scripts).

To reproduce the simulated datasets exactly, the random seed should be retained as specified in the scripts.

## Authors’ contributions

Authors CS, TR, LS and HV have all contributed significantly to the programming and bioinformatics analyses. All authors have contributed to the preparation and writing of this manuscript. All authors have read and approved the final manuscript.

## Acknowledgements

The authors acknowledge the Orion High Performance Computing Center (OHPCC) at the Norwegian University of Life Sciences (NMBU) for providing computational resources that have contributed to the research results reported within this paper (https://orion.nmbu.no).

## Additional files

### File name: Supplementary material 1

File format: .docx

Title: Supplementary material 1

Description: Supplementary material containing Supplementary Table S1 and

Supplementary Figures S1-S6.

### File name: Supplementary material 2

File format: .xls

Title: Supplementary Table S2

Description: Supplementary Table S2. Performance metrics across minimum abundance thresholds and sequencing depths for the VSEARCH-based pipelines. Average values of all evaluated performance metrics for *cluster_size* 99% and UNOISE across minimum abundance thresholds and sequencing depths of 10 000, 50 000, and 100 000 reads. Values are based on 20 replicate datasets for each configuration and are shown for all combinations of species richness and abundance ratio.

### File name: Supplementary material 3

File format: .xls

Title: Supplementary Table S3

Description: Supplementary Table S3. Performance metrics for VSEARCH-based pipelines across minimum abundance thresholds and rarefaction depths. Count-based metrics were derived from empirical environmental sequencing data representing 10 unique sampling sites.

