## Supplementary material 1 for "Benchmarking 16S rRNA gene amplicon analysis in high-diversity microbial communities reveals fundamental trade-offs in clustering and denoising"

**Table S1.** Average number of reads retained after each preprocessing step across simulated datasets. Values represent the average number of reads across 20 replicate datasets for each combination of species richness, abundance ratio, and sequencing depth. DADA2 used reads after trimming and filtering, whereas VSEARCH-based pipelines and Swarm used reads after chimera removal.

| **Number of species** | **Abundance ratio** | **Sequencing depth** | **Average no. of reads after:** | | |
| --- | --- | --- | --- | --- | --- |
|  |  |  | **Trimming and filtering** | **Merging** | **Chimera removal** |
| 100 | 10 | 10 000 | 9 761 | 9 663 | 9 631 |
|  |  | 50 000 | 48 792 | 48 302 | 48 148 |
|  |  | 100 000 | 97 584 | 96 586 | 96 270 |
| 100 | 100 | 10 000 | 9 757 | 9 652 | 9 633 |
|  |  | 50 000 | 48 786 | 48 279 | 48 177 |
|  |  | 100 000 | 97 587 | 96 592 | 96 403 |
| 1 000 | 10 | 10 000 | 9 762 | 9 663 | 9 519 |
|  |  | 50 000 | 48 797 | 48 301 | 47 643 |
|  |  | 100 000 | 97 600 | 96 594 | 95 295 |
| 1 000 | 100 | 10 000 | 9 762 | 9 665 | 9 547 |
|  |  | 50 000 | 48 798 | 48 302 | 47 705 |
|  |  | 100 000 | 97 593 | 96 594 | 95 422 |
| 10 000 | 10 | 10 000 | 9 760 | 9 668 | 9 395 |
|  |  | 50 000 | 48 804 | 48 333 | 46 972 |
|  |  | 100 000 | 97 598 | 96 642 | 94 075 |
| 10 000 | 100 | 10 000 | 9 760 | 9 662 | 9 423 |
|  |  | 50 000 | 48 804 | 48 321 | 47 103 |
|  |  | 100 000 | 97 593 | 96 652 | 94 314 |

**Supplementary Figures**

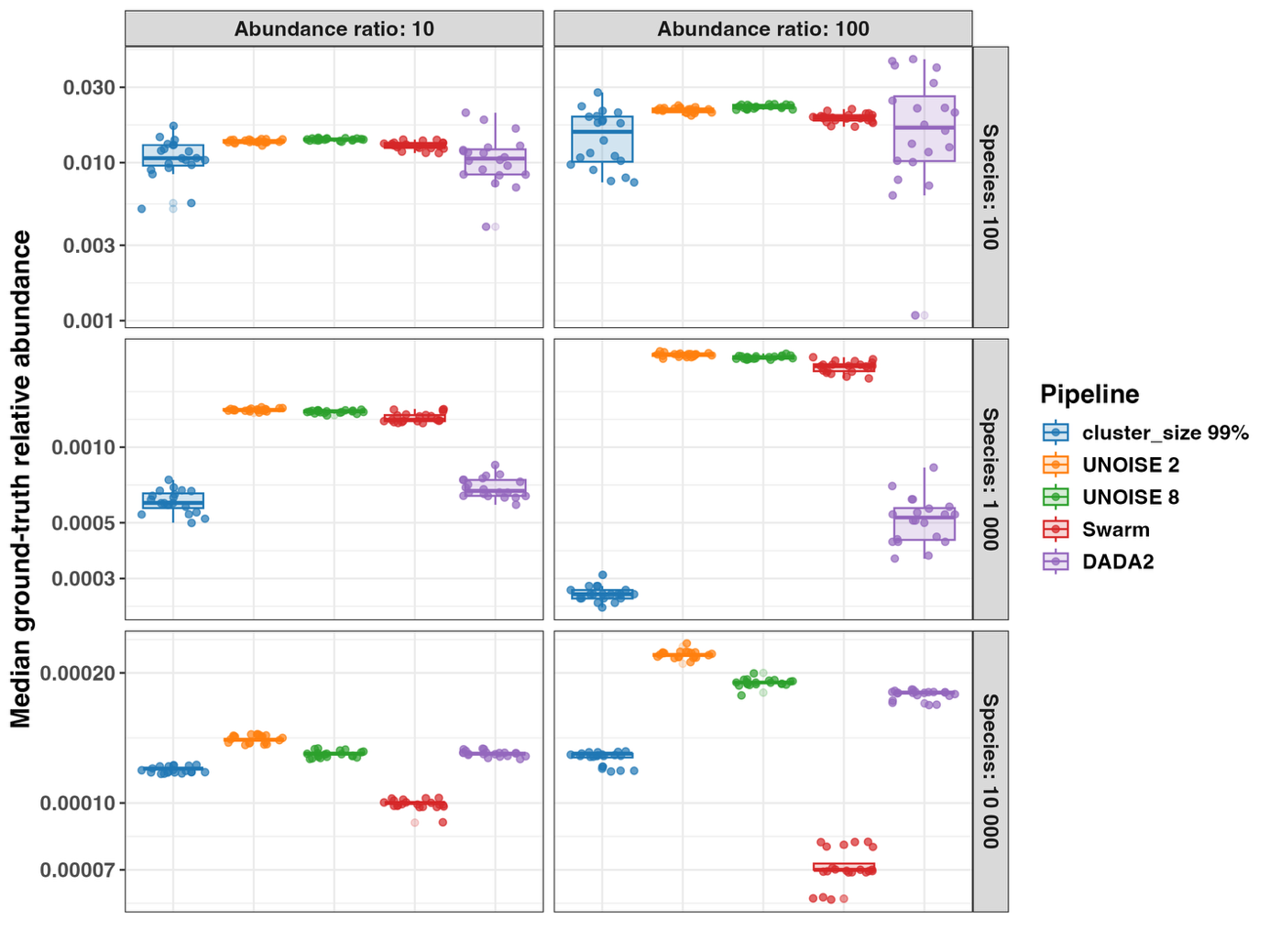
**Figure S1.** Median ground-truth relative abundance of species represented among unclustered reads across clustering and denoising pipelines. Boxplots summarize results across 20 replicate datasets for each combination of species richness and abundance ratio. The y-axis scale varies among species-richness levels; absolute vertical distances should therefore not be compared across rows.

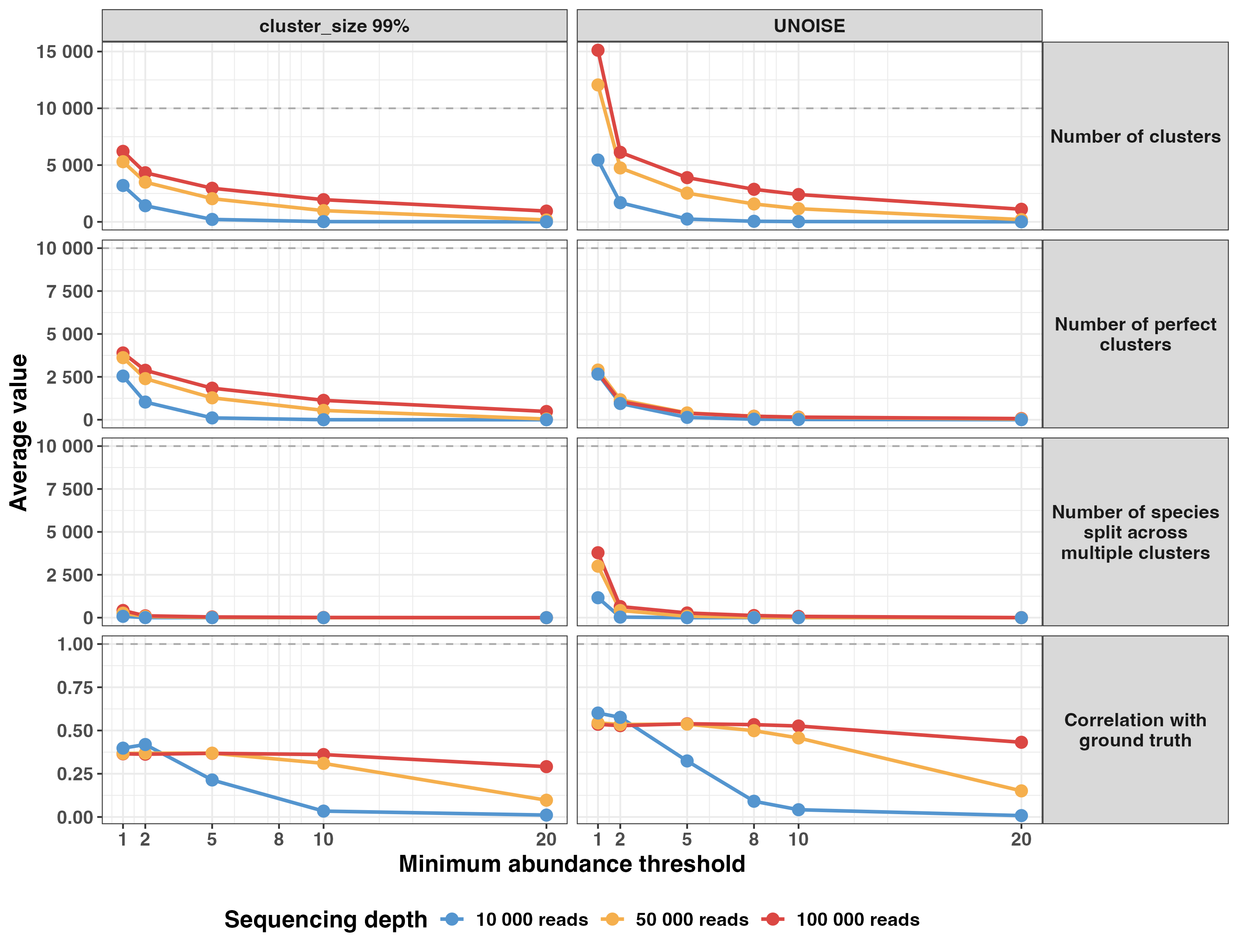
**Figure S2.** Effects of minimum abundance threshold and sequencing depth on the performance of VSEARCH-based clustering and denoising pipelines for the dataset containing 10 000 species and an abundance ratio of 100. Panels compare *cluster_size* at 99% sequence identity and UNOISE across minimum abundance thresholds and sequencing depths of 10 000, 50 000, and 100 000 reads. Points and lines represent average values across 20 replicate datasets for each parameter combination. The horizontal dashed lines indicate the ground-truth species count or the optimal value, where applicable.

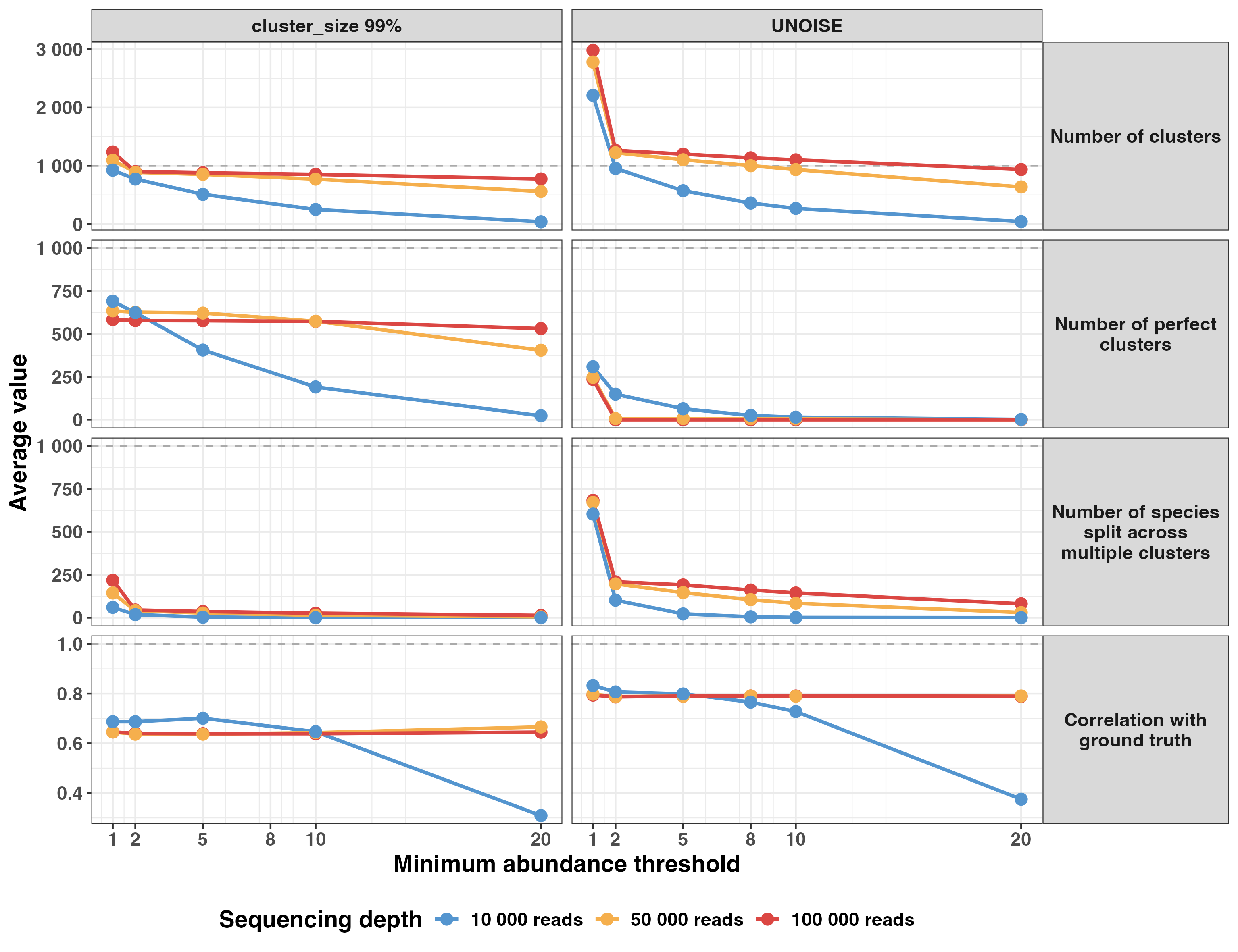
**Figure S3.** Effects of minimum abundance threshold and sequencing depth on the performance of VSEARCH-based clustering and denoising pipelines for the dataset containing 1 000 species and an abundance ratio of 10. Panels compare *cluster_size* at 99% sequence identity and UNOISE across minimum abundance thresholds and sequencing depths of 10 000, 50 000, and 100 000 reads. Points and lines represent average values across 20 replicate datasets for each parameter combination. The horizontal dashed lines indicate the ground-truth species count or the optimal value, where applicable.

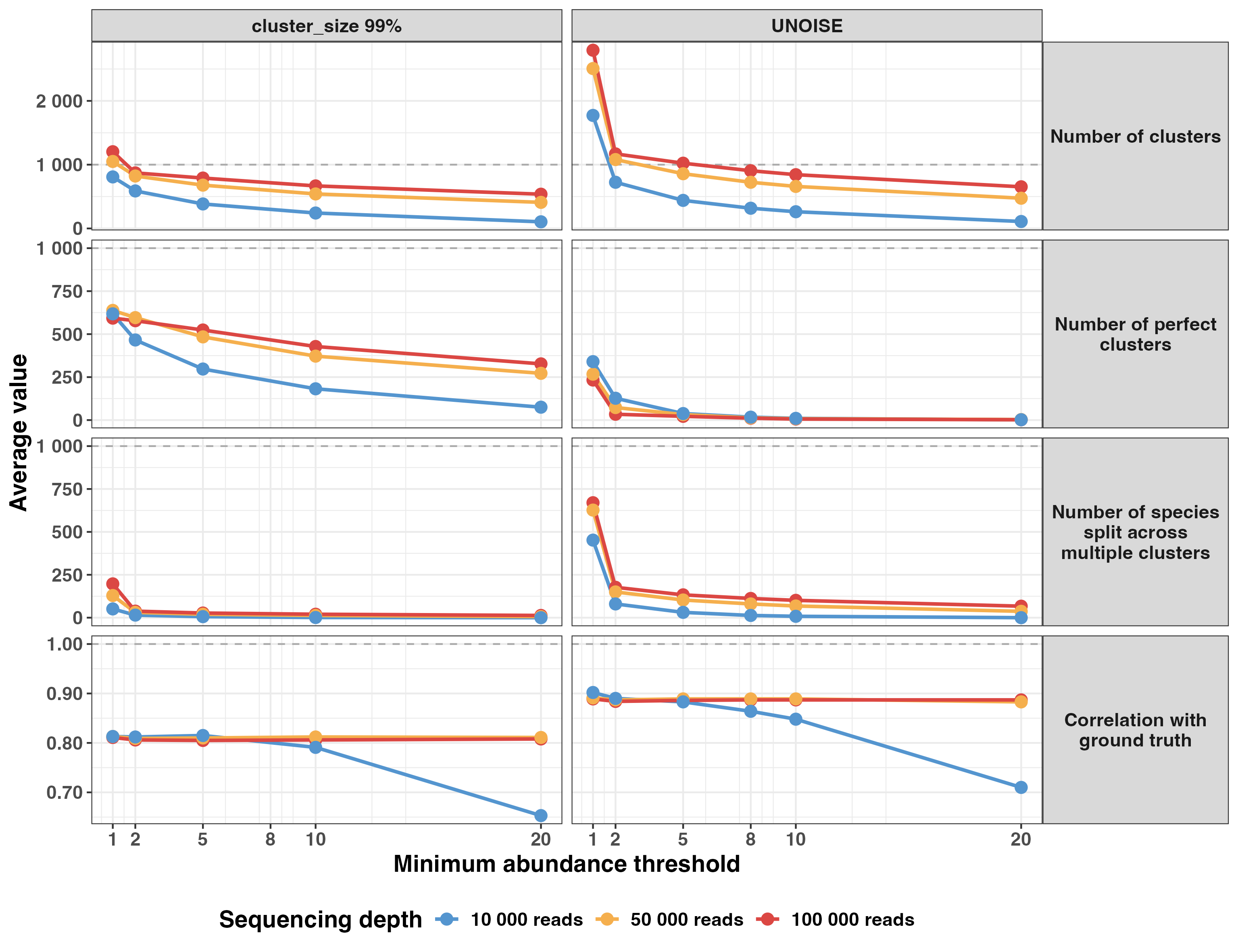
**Figure S4.** Effects of minimum abundance threshold and sequencing depth on the performance of VSEARCH-based clustering and denoising pipelines for the dataset containing 1 000 species and an abundance ratio of 100. Panels compare *cluster_size* at 99% sequence identity and UNOISE across minimum abundance thresholds and sequencing depths of 10 000, 50 000, and 100 000 reads. Points and lines represent average values across 20 replicate datasets for each parameter combination. The horizontal dashed lines indicate the ground-truth species count or the optimal value, where applicable.

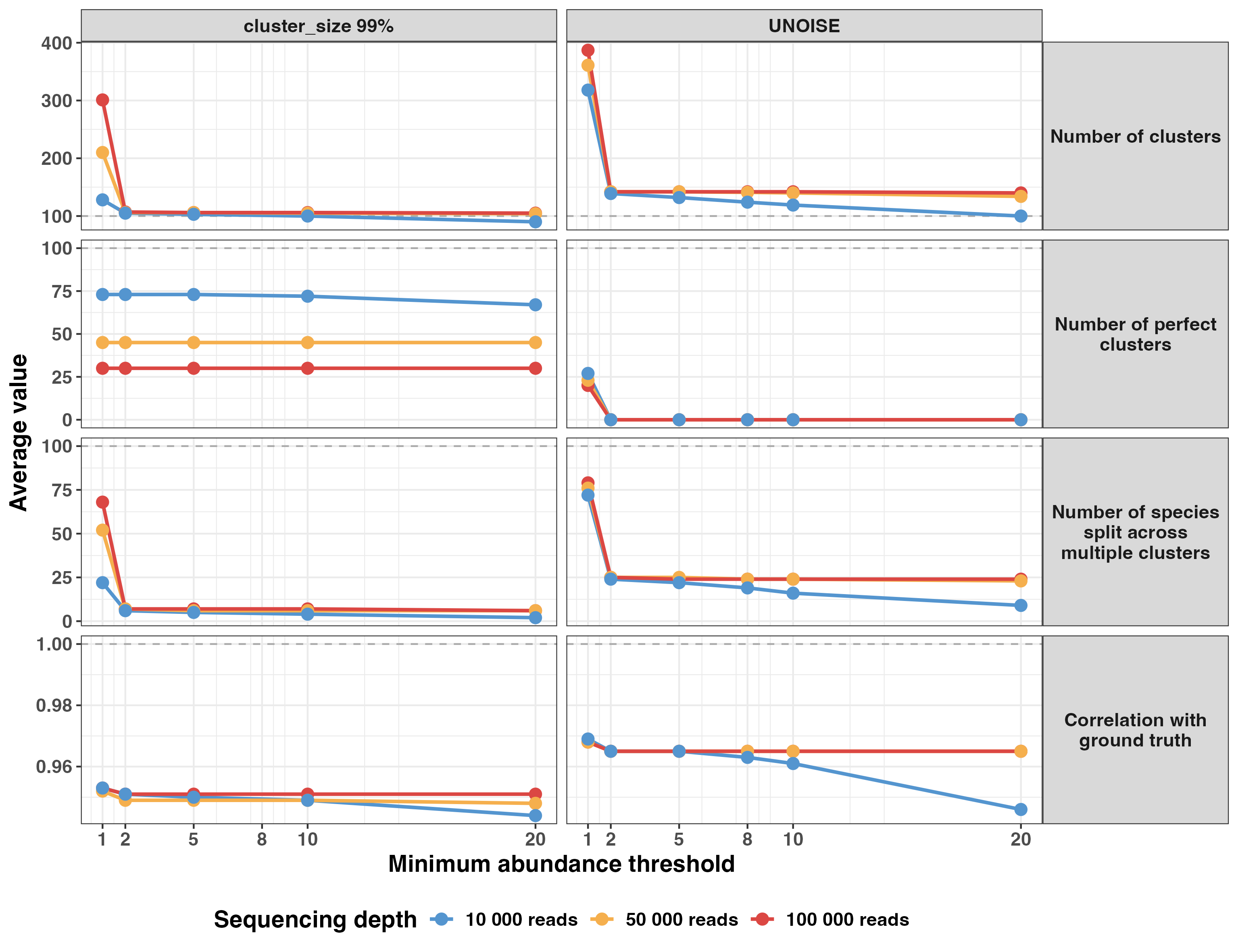

**Figure S5.** Effects of minimum abundance threshold and sequencing depth on the performance of VSEARCH-based clustering and denoising pipelines for the dataset containing 100 species and an abundance ratio of 10. Panels compare *cluster_size* at 99% sequence identity and UNOISE across minimum abundance thresholds and sequencing depths of 10 000, 50 000, and 100 000 reads. Points and lines represent average values across 20 replicate datasets for each parameter combination. The horizontal dashed lines indicate the ground-truth species count or the optimal value, where applicable.

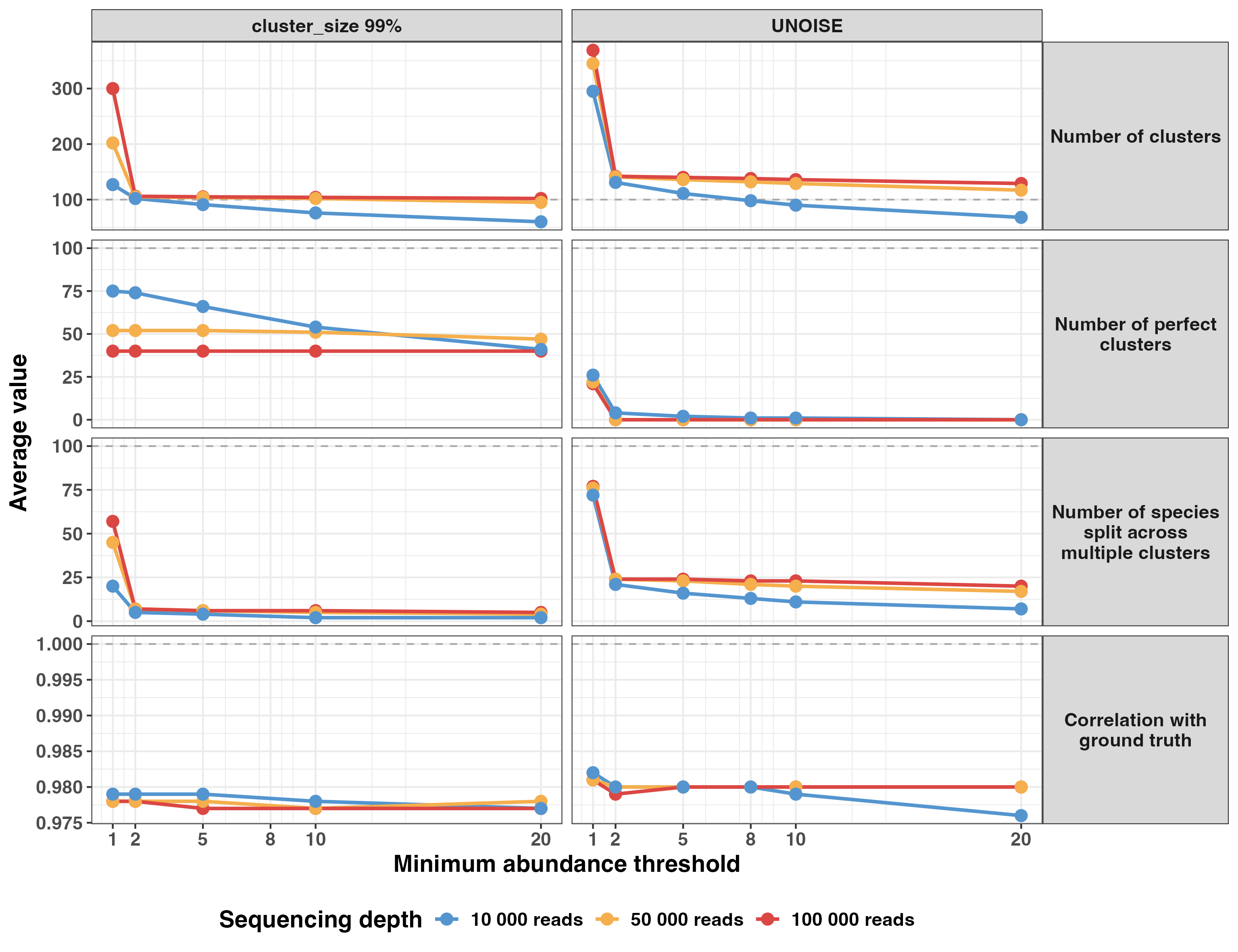

**Figure S6.** Effects of minimum abundance threshold and sequencing depth on the performance of VSEARCH-based clustering and denoising pipelines for the dataset containing 100 species and an abundance ratio of 100. Panels compare *cluster_size* at 99% sequence identity and UNOISE across minimum abundance thresholds and sequencing depths of 10 000, 50 000, and 100 000 reads. Points and lines represent average values across 20 replicate datasets for each parameter combination. The horizontal dashed lines indicate the ground-truth species count or the optimal value, where applicable.
